# Sparse testcrossing for early-stage genomic prediction of general combining ability to increase genetic gain in maize hybrid breeding programs

**DOI:** 10.1101/2025.02.19.639156

**Authors:** David O. González-Diéguez, Gary N. Atlin, Yoseph Beyene, Dagne Wegary, Dorcus C. Gemenet, Christian R. Werner

## Abstract

1

Sparse testcrossing is an effective strategy for increasing both short- and long-term genetic gain in hybrid breeding programs. Maize hybrid breeding programs aim to develop new hybrid varieties by crossing genetically distinct parents from different heterotic pools, exploiting heterosis for improved performance. The programs typically consist of two main components: population improvement and product development. The population improvement component aims to enhance the heterotic pools through reciprocal recurrent selection based on general combining ability (GCA). However, especially in the early stages of testing, evaluating large numbers of hybrid combinations to estimate GCA is impractical due to considerable logistical challenges and costs. Therefore, breeders often evaluate the initial population of selection candidates using only a single tester to narrow down the candidate pool before further evaluation. Using a single tester, however, may not adequately represent the heterotic pool, leading to inaccurate GCA estimates and suboptimal selection decisions. To address this, we propose sparse testcrossing for early-stage testing, where subsets of candidate genotypes are testcrossed with different testers, connected through a genomic relationship matrix. We conducted stochastic simulations to compare various sparse testcrossing designs with a conventional testcross strategy using a single tester over 15 cycles of reciprocal recurrent genomic selection. Our results show that using 3-5 testers, sparsely distributed among full-sibs, sparse testcrossing offers breeders a practical balance between simple testcross designs, resource efficiency, and increased prediction accuracy for GCA, ultimately resulting in increased rates of genetic gain.

**Key message:** Sparse testcrossing with 3-5 testers enhances genetic gain in hybrid breeding programs, offering a practical balance of simple testcross designs, resource efficiency, and increased prediction accuracy for general combining ability.

## 2 INTRODUCTION

Sparse testcrossing is an effective strategy for increasing both short- and long-term genetic gain in hybrid breeding programs. By using 3–5 testers, sparsely distributed among full-sibs, it offers breeders a practical balance between simple testcross designs, resource efficiency, and increased prediction accuracy for general combining ability (GCA) compared to conventional testcross strategies using a single tester. As such, sparse testcrossing lays the foundation for rapid recycling of parents at the earliest stage of agronomic testing to minimize breeding cycle time and maximize genetic gain per unit of time.

Maize hybrid breeding programs aim to develop high-performing hybrid varieties by crossing genetically distinct parental genotypes, typically derived from different populations referred to as heterotic pools. In this way, hybrid breeding exploits heterosis (hybrid vigor), which results from the combination of complementary beneficial haplotypes from these heterotic pools, leading to improved performance in the hybrids compared to their parents (Melchinger and Gumber 1998). Hybrids can be produced through various methods, including single cross (crossing two inbred lines), three-way cross (crossing a single cross with a third inbred line), or double cross (crossing two single crosses) (Bernardo 2020). While many commercial programs with a long history of hybrid maize breeding typically focus on single-cross hybrids, public sector programs and small to medium-sized seed companies often use three-way crosses, with the maternal component being a single-cross hybrid, to compensate for low seed set caused by inbreeding depression.

Maize hybrid breeding programs typically consist of two key components: (1) population improvement and (2) product development. The population improvement component focuses on enhancing the genetic composition of the heterotic pools for both per se performance and complementarity between the pools, thereby creating superior hybrid parents.

Population improvement is achieved through reciprocal recurrent selection of genotypes, such as inbred lines, based on GCA (Comstock et al. 1949). The GCA predicts the average performance of a genotype when randomly crossed with genotypes from the complementary heterotic pool (Hallauer et al. 2010). It reflects both additive genetic effects and, to some extent, non-additive effects that arise from complementary haploblock interactions and serves as a predictor of a parent’s ability to produce high-performing hybrids when crossed with other genotypes from a complementary heterotic pool (Bernardo 2020). Following population improvement, the product development component then focuses on identifying the best-performing hybrid combinations among all potential reciprocal crosses between parental genotypes from the improved heterotic pools.

Ideally, GCA is estimated in a complete factorial mating design, which evaluates all possible hybrid combinations between heterotic pools (Comstock and Robinson 1948; Fritsche-Neto et al. 2018). However, the implementation of such a mating design at early stages of testing is often impractical due to the large number of hybrid combinations and the associated logistical challenges and costs. To address this, maize breeders typically use one or a few testers, selected to represent the complementary heterotic pool (Hallauer et al. 2010).

In many public maize hybrid breeding programs, it is common practice to employ a two-stage selection approach before parents are recycled. In stage 1, the initial population of candidate genotypes is typically evaluated using a single tester, providing a preliminary assessment to narrow down the candidate pool. In stage 2, the pre-selected subset of genotypes is further evaluated using additional testers – usually 3 to 5 – to enhance GCA accuracy, as more testers better capture the genetic diversity and haploblock frequencies in the complementary heterotic pool. However, this approach comes at the cost of extending the breeding cycle time and slowing genetic gain. Additionally, pre-selection with a single tester may not adequately represent the complementary heterotic pool, leading to inaccurate GCA estimates and suboptimal advancement of candidates to stage 2 testing.

Genomic selection offers great potential to increase GCA accuracy in stage 1 testing, thereby increasing genetic gain by enabling earlier parent selection and more efficient product advancement. By leveraging haplotype blocks as the primary units of evaluation rather than individual genotypes (Bingham 1998; Meuwissen et al. 2001), genomic selection enhances prediction accuracy and can reduce the breeding cycle, thereby accelerating genetic gain (Crossa et al. 2017). Genomic selection has already proven to be a valuable tool to improve maize hybrid breeding, particularly for predicting hybrid performance (Technow et al. 2014; Kadam et al. 2016; Fritsche-Neto et al. 2018; Seye et al. 2020) and predicting GCA for parent recycling (Melchinger and Frisch 2023; de Jong et al. 2023; Lorenzi et al. 2024). Studies focused on genomic prediction of GCA suggest that incomplete factorial designs improve GCA prediction accuracy and genetic gain compared to single-tester designs (Melchinger and Frisch 2023; Lorenzi et al. 2024). However, incomplete factorial designs require labor-intensive hand pollination, which is costly and challenging, particularly when synchronizing flowering times. In contrast, conventional testcross approaches with a single tester often use open pollination for producing testcrosses, simplifying logistics, but resulting in low GCA accuracy (Fritsche-Neto et al. 2018). Therefore, novel strategies are needed to balance practicality, simple testcrossing designs, and prediction accuracy.

In this study, we propose sparse testcrossing for early-stage testing using multiple testers in sparse designs, leveraging genomic relationships to connect candidate genotypes. While each candidate is crossed with only a single tester, related candidate genotypes with shared haplotypes are tested with different testers, allowing for a broader sampling of haplotypes within the complementary heterotic (tester’s) pool. As a result, the genomic predicted values of the haplotypes in a single genotype reflect their average performance across multiple testcrosses rather than just one. We hypothesize that this strategy enhances GCA prediction accuracy, enabling early-stage parent selection without increasing resource demands compared to a conventional single-tester testcross design. This strategy builds on the idea of sparse testing (Endelman et al. 2014; Werner et al. 2025), treating testers as distinct “testing environments”, with subsets of candidate genotypes testcrossed with different testers and relying on genomic relationships to maintain connectivity among selection candidates. To assess its effectiveness, we conducted stochastic simulations comparing ten sparse testcrossing designs against a conventional testcross design with a single tester across 15 cycles of reciprocal recurrent genomic selection, representing the population improvement component of a rapid cycle genomic selection hybrid breeding program.

## 3 MATERIALS AND METHODS

Stochastic simulations were conducted to compare ten different sparse testcrossing designs for genomic prediction of general combining ability (GCA) to facilitate early-stage parent selection in hybrid breeding programs. The sparse testcrossing designs were evaluated for prediction accuracy and hybrid genetic gain over 15 cycles of recurrent reciprocal genomic selection and compared to a conventional early-stage testcross strategy with a single tester. Comparisons were based on a single quantitative trait, such as yield, under three different dominance degrees.

The material and methods are subdivided into five sections:

i) Simulation of the founder population and heterotic pool formation
ii) Description of the sparse testcrossing designs
iii) Simulation of 15 cycles of recurrent reciprocal genomic selection
iv) Genomic prediction models
v) Evaluation of sparse testcrossing designs for GCA prediction accuracy and genetic gain

### 3.1 Simulation of the founder population and heterotic pool formation

#### 3.1.1 Genome simulation

Whole-genome sequences were simulated for a founder population of 80 heterozygous genotypes. To roughly mimic the maize genome, each founder genome consisted of 10 chromosome pairs with a physical length of 2 x 10^8^ base pairs (bp) and a genetic length of 200 centimorgans (cM), resulting in a total physical length of 2 Gbp and a genetic length of 2000 cM. Whole-genome sequences were generated using the Markovian coalescent simulator (Chen et al. 2009) in AlphaSimR (Gaynor et al., 2021), deploying AlphaSimR’s maize evolutionary history default settings. Recombination rate was derived as the ratio between genetic length and physical length (i.e., 2 Morgan/2×10^8^ bp = 10^-8^ recombinations per base pair), and mutation rate was set to 1.25×10^–8^ per base pair. The effective population size (Ne) at the end of the coalescent process was set to 100. The main purpose of the coalescence process was to generate linkage disequilibrium in the founder population.

A set of 500 biallelic quantitative trait nucleotides (QTN) and 500 single nucleotide polymorphisms (SNP) were randomly sampled along each chromosome to simulate a quantitative trait controlled by 5,000 QTN and a SNP marker array with 5,000 markers. Founder genotypes were formed by randomly sampling 10 chromosome pairs per genotype.

#### 3.1.2 Simulation of genetic values

Genetic values for a single quantitative trait were simulated by summing biological additive genetic effects and dominance effects at the 5,000 QTN. Additive genetic effects (*a*) were randomly sampled from a standard normal distribution and scaled to obtain an additive genetic variance in the founder population of 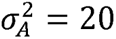, following a study by De Jong et al., 2023 on the comparison of genomic prediction models for GCA in early stages of hybrid breeding programs.

Dominance effects (*d*) were calculated by multiplying the absolute biological additive effect of a QTN (*ai*) by a locus-specific dominance degree (*δ_i_*), that is *d_i_* = *δ_i_* × |*a_i_*| Dominance degrees were sampled from a normal distribution with mean dominance coefficient *μ_δ_* and variance 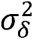, that is 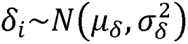 Three levels of average dominance degrees, 0.1, 0.5 and 0.9, were used to simulate overall positive directional dominance. The dominance degree variance 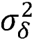 was set to 0.2. As reference, the ratios of SCA to GCA variances in the first cycle were 5%, 23% and 50% for low (*μ_δ_*=0.1), medium (*μ_δ_*=0.5) and high (*μ_δ_*=0.9) dominance degrees, respectively.

#### 3.1.3 Formation of initial heterotic pools

The 80 founder genotypes were converted into homozygous inbred lines and randomly allocated to one of two gene pools consisting of 40 inbred lines each (Fig. 1A). The two gene pools served as separate male and female heterotic pools. To create a baseline family structure within heterotic pools as a basis for genomic prediction, three generations of random crossing and selection were conducted. At first, 60 biparental crosses were sampled from a half diallel without selfing to produce F_1_ genotypes, ensuring that each of the 40 inbred lines was used in at least one cross. Then, 20 fully homozygous inbred lines were created from each F_1_, resulting in a total of 1,200 inbred lines per heterotic pool. Among those inbred lines, 40 lines were randomly selected as new parents. This procedure was repeated for two more generations, resulting in a slight differentiation between pools (Fig. 1B). To generate fully homozygous inbred lines, the *makeDH()* function in AlphaSimR was used, which generates double haploid (DH) lines.

**Fig. 1.**
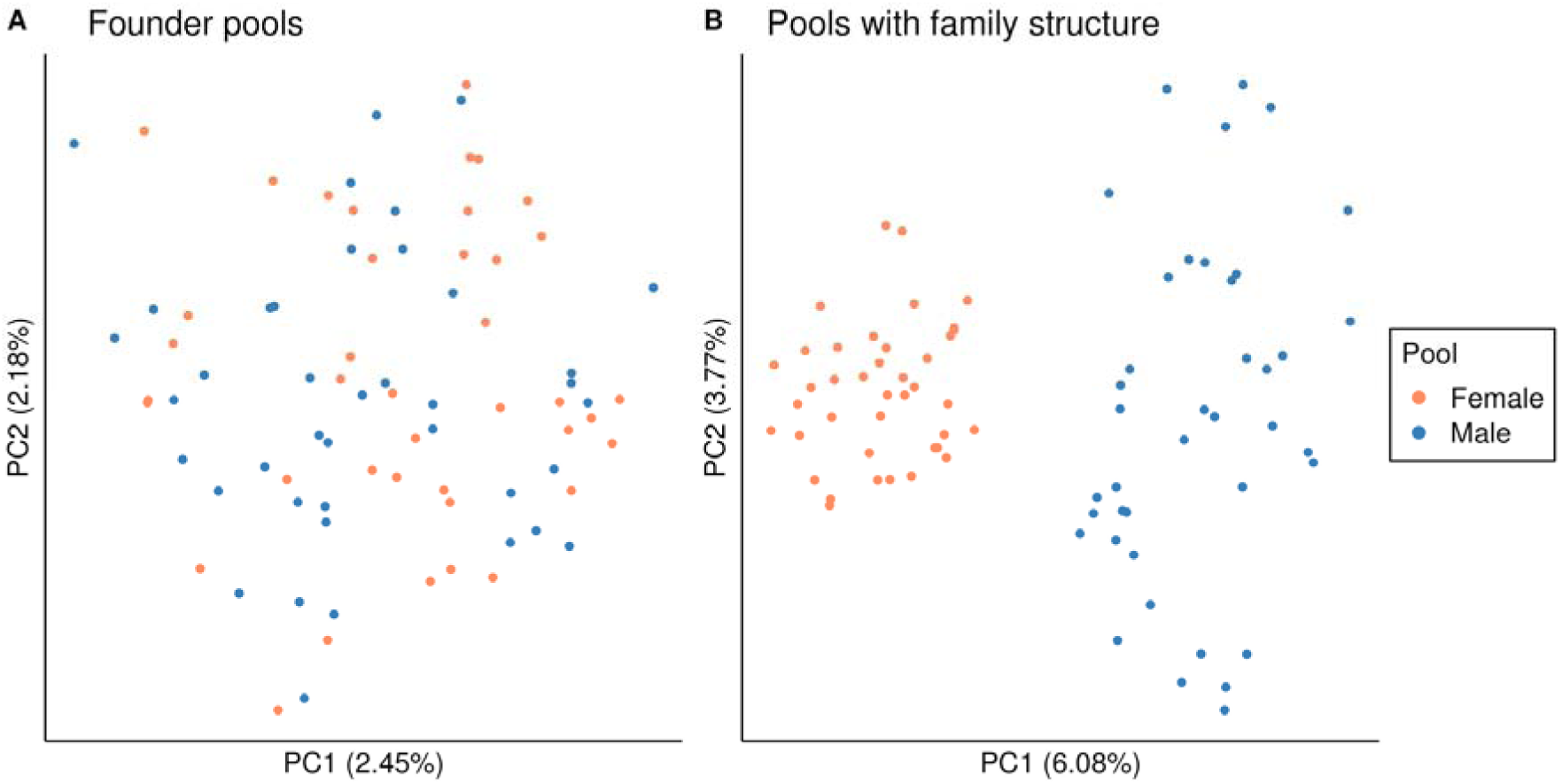
The first two principal components from a principal component analysis (PCA) performed on the genomic relationship matrix of the inbred lines of the founder pools in the baseline breeding program, before (A) and after (B) generating baseline family structure within pools. The first principal component is represented on the X-axis, while the second principal component is plotted on the Y-axis. Colors orange and blue represent female and male heterotic pools, respectively. The proportion of variance explained by the first and second principal components (PC) is indicated in brackets. Results correspond to a single, randomly selected simulation replication of the baseline breeding program with medium dominance degree

### 3.2 Description of the sparse testcrossing designs

We compared ten sparse testcrossing designs in which each tester from one heterotic pool was crossed with an equally sized subset of inbred lines from the other heterotic pool. Subsets of candidate lines were created at random, allowing full-sibs to be crossed with different testers. The ten sparse testcrossing designs used sets of 2, 3, 4, 5, 10, 20, 50, 100, 600, and 1,200 testers, respectively, to generate 1,200 testcrosses per heterotic pool (Table 1). A conventional early-stage testcross design with a single tester crossed to all 1,200 inbred lines was used as a benchmark. Randomly selected inbred lines served as testers, and a new set of testers was sampled at each crossing cycle.

**Table 1.** Number of testers and testcrosses per tester used in the conventional early-stage testcross design with a single-tester (benchmark) and the ten sparse testcrossing designs.

| # Testers | 1 | 2 | 3 | 4 | 5 | 10 | 20 | 50 | 100 | 600 | 1200 |
| --- | --- | --- | --- | --- | --- | --- | --- | --- | --- | --- | --- |
| # Crosses per Tester | 1,200 | 600 | 400 | 300 | 240 | 120 | 60 | 24 | 12 | 2 | 1 |

### 3.3 Simulation of 15 cycles of recurrent reciprocal genomic selection

To compare the ten sparse testcrossing designs and the conventional early-stage testcross design with a single-tester for genomic prediction accuracy of GCA and hybrid genetic gain, we simulated 15 cycles of recurrent reciprocal genomic selection, representing the population improvement component of a rapid cycle genomic selection hybrid breeding program with inbred lines. The basic structure of the recurrent reciprocal genomic selection scheme is shown in Fig. 2, and hereinafter described for one heterotic pool.

**Fig. 2.**
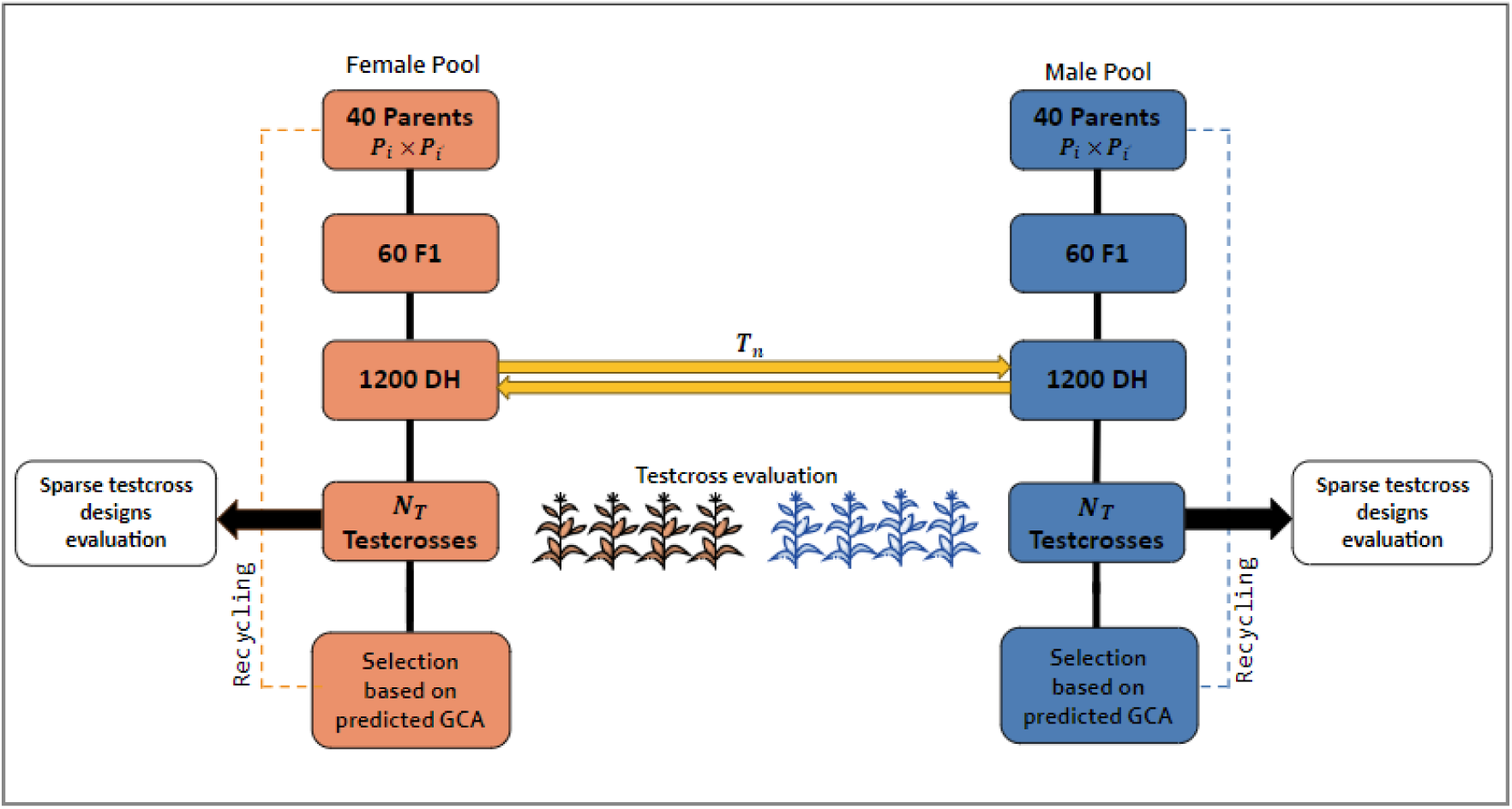
Schematic representation of the rapid cycling double haploid hybrid breeding scheme. A total of 15 cycles of recurrent reciprocal genomic selection were simulated. represents the number of testcrosses. represents the number of testers. Maize plant icons were sourced from Flaticon.com

At each cycle, 60 biparental crosses were sampled from a half diallel without selfing between the 40 inbred lines to generate F1 genotypes. Then, from each F1, 20 fully homozygous inbred lines were derived, resulting in a total of 1,200 inbred lines. These inbred lines were crossed with testers from the other heterotic pool to produce testcrosses. Testcross phenotypes were generated by adding a random error to the testcross genotypic values. The random error was sampled from a normal distribution with mean zero and the error variance set to obtain a broad-sense heritability of *H^2^* = 0.3 in the hypothetical population of all 40 x 40 = 1600 possible hybrids between the initial male and female heterotic pools, i.e., , with being the genetic variance of the hypothetical hybrid population. The error variance was kept constant across all 15 cycles of recurrent reciprocal selection. Genomic predicted GCA was then calculated by regressing testcross phenotypic performance on the inbred line SNP allele dosages, as described in the following section, and the 40 inbred lines with the highest GCA were recycled as new crossing parents.

### 3.4 Genomic prediction models

Depending on the testcross design, two different models were used for genomic prediction of GCA. In the conventional early-stage testcross design with a single-tester, we used a model which calculated average effects of SNP marker alleles for testcross performance specific to each heterotic pool (Eq. 1). This is because only one tester was used and the SCA effects could not be separated from GCA effect. This model is hereafter referred to as “Testcross Model”, following the terminology used by De Jong et al. (2023).

In the sparse testcrossing designs with more than one tester, we used a model which jointly calculated heterotic pool-specific biological additive allele effects (i.e., for the inbred lines and the testers), as well as biological dominance effects for heterozygous SNP marker loci in the testcrosses (Eq. 2). The use of multiple testers reduces the influence of SCA effects on the estimation of the GCA effects, and this model allows better delineation of the GCA and SCA effects while accounting for pool-specific LD between QTL and markers, as reported by De Jong et al. (2023). To predict the GCA of inbred lines, the *setEBV()* function in AlphaSimR was employed. As described by De Jong et al. (2023), this function utilizes both biological additive and dominance effects to calculate the average effect of an allele for each SNP marker following Eq. 3. These average effects are then used to predict an inbred line’s GCA. This model accounts for directional dominance and is hereafter referred to as “Hybrid Pool Specific Additive Effects + Dominance Model”, following the terminology used by De Jong et al. (2023).

Both models were fitted using the ridge regression best unilinear prediction (RRBLUP) and REML variance component estimation framework in AlphaSimR.

#### 3.4.1 Testcross Model

The Testcross Model used for genomic prediction of GCA in the conventional early-stage testcross design with a single-tester is as follows:

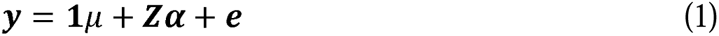

where ***y*** is the *n* × 1 vector of testcross phenotypes with *n* being the number of testcrosses, 1 is an *n* × 1 vector of ones, is the intercept; ***Z*** is the *n* × *m* matrix of inbred line genotypes, with *m* being the number of SNP markers coded as 0 and 2 for genotypes aa and AA; ***α*** is an *m* × 1 vector of average effects of SNP marker alleles; and *e* is the *n* × 1 vector of residuals. The model was fitted using the AlphaSimR function *RRBLUP()*. The of an inbred line was predicted by summing the products of the marker effects and the inbred line’s corresponding marker genotypes.

#### 3.4.2 Hybrid Pool Specific Additive Effects + Dominance Model

The Hybrid Pool Specific Additive Effects + Dominance Model, used for genomic prediction of GCA in the sparse testcrossing designs, fits additive effects for each heterotic pool and jointly estimates dominance effects across both heterotic pools. Additionally, the model accounts for directional dominance, following the methodology described by Xiang et al. (2016). A detailed description of this model is provided by De Jong et al. (2023) and is presented as follows:

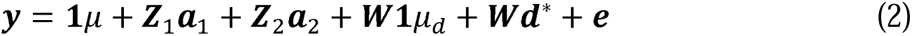

where ***y*** is the *n* × 1 vector of testcross phenotypes with *n* being the number of testcrosses, ***Z***_1_ is the *n* × *m* genotype matrix of inbred lines from one heterotic pool with *m* being the number of SNP markers coded as 0 and 1 for genotypes aa and AA; ***Z***_2_ is the *n* × *m* genotype matrix of the DH lines used as testers from the other heterotic pool, with entries coded as in ***Z***_1_. *W* is an *n* × *m* heterozygosity matrix coded as 0, 1, and 0 for testcross genotypes AA, Aa, and aa, respectively; ***W***1 is an *n* × 1 vector containing individual heterozygosity for each testcross hybrid; ***a***_1_ and ***a***_2_ are the *m* × 1 vectors of biological additive effects for inbred lines and testers, respectively; *p_d_* is an average of dominance effects for a single heterozygous locus and ***W***1*μ_d_* results in an average of dominance effects or genomic individual heterosis for each testcross hybrid; ***d****\** is an *m* × 1 vector of dominance effects not captured by ***W***1*μ_d_ (i. e., d* = d* − *μ_d_ )W* with *E (d\**) = 0; and *e* is the *n* × 1 vector of residuals. This model was fitted using the AlphaSimR function *RRBLUP_SCA()*.

According to De Jong et al. (2023), pool-specific biological additive effects (*a*) and dominance *(d)* effects are used in AlphaSimR to calculate the average testcross effects of an allele substitution for each SNP marker in heterotic pool 1 and 2 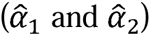, respectively:

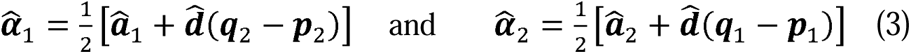

with *p*_1_ and *q*_1_ = 1 − *p_1_* being the SNP marker allele frequencies in pool 1; and *p_2_* and *q_2_* being the SNP markers allele frequencies in pool 2. Similar expressions were presented by Gonzalez-Dieguez et al. (2021). The GCA of an inbred line was then predicted by summing the products of the marker effects and the inbred line’s corresponding marker genotypes.

### 3.5 Evaluation of sparse testcrossing designs for GCA prediction accuracy and hybrid genetic gain

#### 3.5.1 Evaluation of GCA prediction accuracy

To ensure a fair comparison of the ten sparse testcrossing designs and the conventional early-stage testcross design with a single tester, the accuracy of genomic predicted GCA was measured at the parent selection stage of the same baseline recurrent reciprocal genomic selection breeding program. This was done to prevent the prediction accuracies from being affected by different genetic variance trajectories over crossing cycles, which is observed when each testcross design is evaluated in a separate breeding program simulation (Suppl. Fig. S10).

The baseline breeding program mimics the population improvement component of a conventional reciprocal recurrent genomic selection hybrid breeding program (Fig. 2), in which inbred lines are usually crossed to and evaluated with several testers before new parents are identified. Therefore, each inbred line was crossed with the same three testers, resulting in a total of 3 x 1,200 = 3,600 testcrosses per heterotic pool. Note that this corresponds to three times the testcross resources used in the sparse testcrossing designs. Also, the average testcross performance of inbred lines was used for genomic selection in the baseline breeding program. The process of selecting parents based on genomic predicted GCA is expected to generate divergence between heterotic pools over crossing cycles, which will enable the examination of the effect of genetic distance between pools on genomic prediction accuracy of GCA.

Genomic prediction accuracy of GCA of the ten sparse testcrossing designs and the single-tester design was measured at cycles 1, 2, 3, 5, 10, and 15 of the baseline breeding program. Prediction accuracy was calculated as the Pearson correlation coefficient between genomic predicted GCA and true GCA. The true GCA was calculated as the mean genetic value of an inbred line from one heterotic pool crossed with all inbred lines from the other heterotic pool. Phenotypic prediction accuracy of GCA served as a benchmark.

#### 3.5.2 Evaluation of genetic gain

To compare the ten sparse testcrossing designs and the conventional early-stage testcross design with a single-tester for their effect on hybrid genetic gain, 15 cycles of recurrent reciprocal genomic selection (Fig. 2) were simulated for each testcross strategy. Hybrid genetic gain was measured as the mean total genotypic value of all 1,440,000 potential hybrids (full factorial) between the two heterotic pools. Genetic gain was expressed relative to the mean genetic value of hybrids formed between initial heterotic pools. While genomic prediction accuracies of GCA were also measured for each testcross strategy (Suppl. Fig. S12), a direct comparison between strategies is difficult since prediction accuracies were affected by different genetic variance trajectories over crossing cycles (Suppl. Fig. S10).

#### 3.5.3 Divergence of heterotic pools, genetic variances and diversity

The divergence of heterotic pools, genetic variance components and genetic diversity were monitored over crossing cycles in the baseline breeding program and within each recurrent reciprocal genomic selection program for each testcrossing strategy.

Heterotic pool divergence was assessed using three metrics: (1) Nei’s minimum genetic distance between the pools, (2) the average heterosis observed between the two pools, and (3) the fraction of fixed alleles in the two pools. The Nei’s minimum genetic distance was calculated as 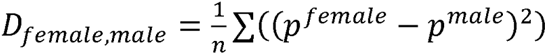 (Nei 1987), where *n* is the number of polymorphic QTN, and *p^female^* and *p^male^* are the allele frequencies in the female and male heterotic pool, respectively. Mid-parent heterosis, defined as the deviation from the mean of the two parental populations, was calculated as described in Falconer (1981) (p. 257 Eq. 14.8), using the equation 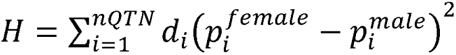. The fraction of fixed alleles was calculated relative to the number of QTNs. Quantitative trait nucleotides with allele frequencies of 0 in one pool and 1 in the other pool were considered fixed for opposite alleles, while QTNs with allele frequency of 0 or 1 in the two pools were considered fixed for the same allele.

Genetic variance components were analyzed by monitoring the total genetic variance of hybrids, the GCA and SCA variances. The total genetic variance of hybrids was calculated from the true total genetic values of hybrids. The GCA variance was calculated from the true GCA values of inbred lines. The SCA variance (or dominance deviation variance, given that epistatic effects were not simulated) was calculated by subtracting the GCA variances from the total genetic variance of hybrids.

For genetic diversity, the number of families of selected parents and the Jaccard similarity coefficient (Jaccard 1908) within each pool were calculated. The Jaccard similarity coefficient was calculated using SNP maker genotypes of selected inbred parents within each pool. A similarity matrix was generated using the *vegdist()* function from the *vegan* R package (Oksanen et al. 2001). The average value of the lower triangular portion of this matrix was then used as the final similarity metric. Results of the number of families and Jaccard similarity coefficient were averaged across both female and male heterotic pools.

## 4 RESULTS

Our results show that sparse testcrossing can improve genomic prediction accuracy of general combining ability (GCA) compared to a conventional early-stage testcross strategy using a single tester, thereby facilitating increased rates of hybrid genetic gain. Both genomic prediction accuracy of GCA and hybrid genetic gain increased with the number of testers, reaching a plateau around 10 testers. Sparse testcrossing improved prediction accuracy and genetic gain most during the early breeding cycles, while these advantages decreased as the genetic distance between pools increased. Furthermore, the benefits of sparse testcrossing were more pronounced when the degree of dominance was high but diminished as the degree of dominance decreased. Since similar results were observed in both heterotic pools, we only present the results for one of the two pools, focusing on the scenario with a medium dominance degree. The low and high dominance degree results are only addressed when relevant to the interpretation of the results, and otherwise can be found in the supplementary material.

### 4.1 Divergence of heterotic pools, genetic variances and diversity in the baseline breeding program

Selection of parents based on GCA resulted in the divergence of heterotic pools over time. This is shown in Fig. 3, which presents the first two principal components from a principal component analysis (PCA) performed on the genomic relationship matrix of the candidate inbred lines at cycles 1, 2, 3, 5, 10, and 15. For example, at cycle 1 a low population structure is observed, as the first principal component account only 6.16% of the total variance (Fig. 3A). By cycle 15, this proportion had increased to 92.94%, with the first principal component reflecting a large between-pool variance (Fig. 3F). Figure 3 illustrates the effectiveness of parent selection based on GCA in quickly creating distinct heterotic pools and enhancing hybrid performance, a hallmark of an efficient hybrid breeding program. The data shown in Fig. 3 comes from a single, randomly selected simulation replication of the baseline breeding program with a medium dominance degree. Similar patterns of heterotic pool divergence over time were also observed for the baseline breeding program with low and high degrees of dominance (Suppl. Figs. S1 and S2).

**Fig. 3.**
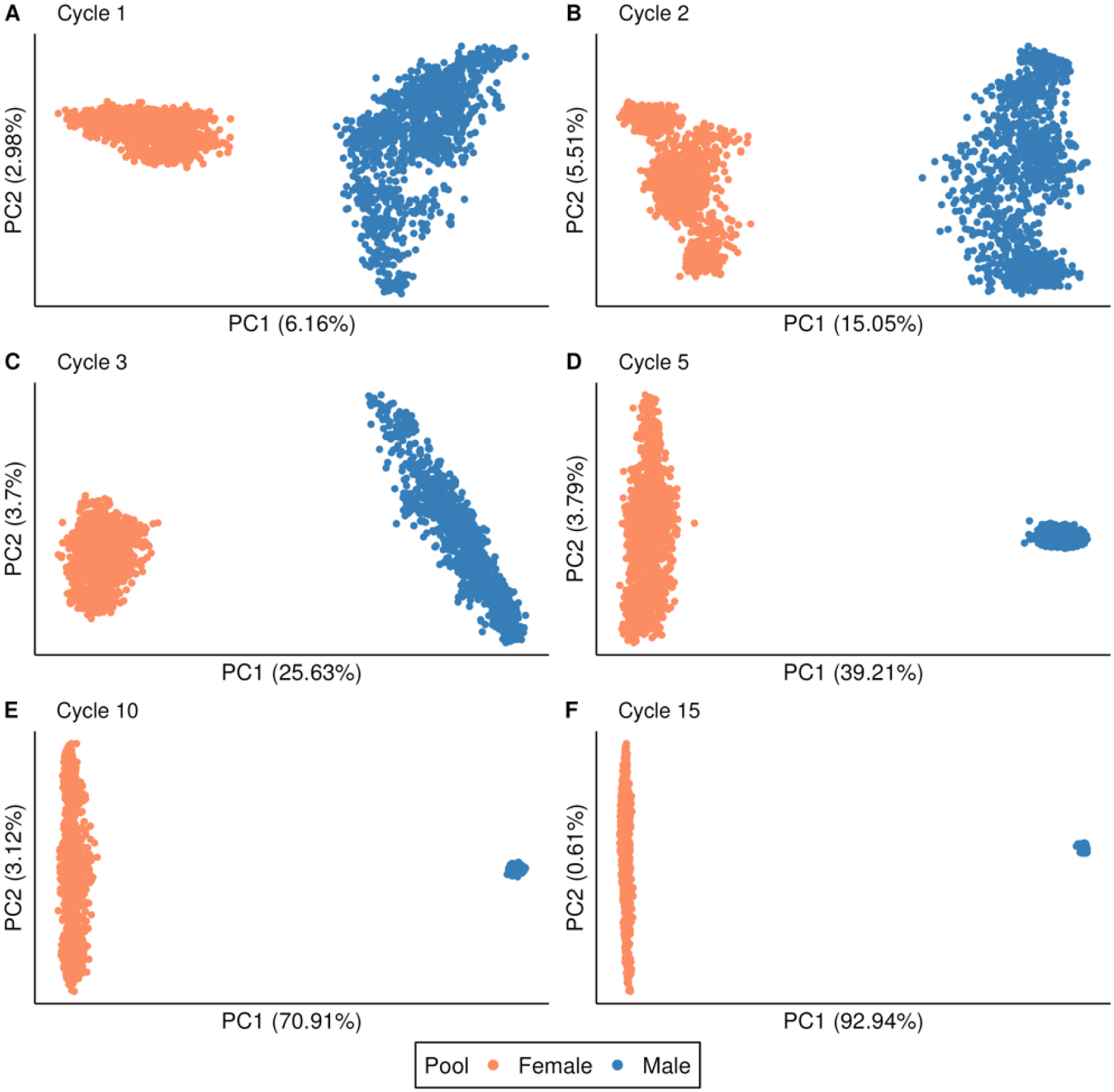
The first two principal components from a principal component analysis (PCA) are shown, based on the genomic relationship matrix of candidate inbred lines at cycles 1, 2, 3, 5, 10, and 15 of the baseline breeding program. The first principal component is plotted on the X-axis, while the second principal component is on the Y-axis. The colors orange and blue represent female and male heterotic pools, respectively. The proportion of variance explained by the first and second principal components (PC) is indicated in brackets. These results are derived from a single, randomly selected simulation replication of the baseline breeding program with medium degree of dominance

The divergence of the two heterotic pools over time led to an increase in genetic distance between pools, an increase in heterosis, and a greater fraction of fixed alleles in the two pools. This is illustrated in Fig. 4, which shows the mean genetic distance (Fig. 4A), mean heterosis (Fig. 4B), and mean fraction of fixed (same or opposite) alleles in the two pools (Fig. 4C) across all 100 simulation replications of the baseline breeding program with low, medium and high degree of dominance, respectively. Additionally, as consequences of the selection process, the total genetic variance of hybrids, as well as the GCA and SCA variances, declined over time. This is shown in Fig. 5, which plots the mean total genetic variance of hybrids, the mean GCA variance and mean SCA variance across all 100 simulation replications of the baseline breeding program with high (Fig. 5A), medium (Fig. 5B) and low (Fig. 5C) dominance degrees, respectively. As expected, after 15 cycles of crossing and selection, genetic distance and heterosis between the two pools were higher when the dominance degree was high, resulting in a more pronounced depletion of GCA and SCA variances. Additionally, the fraction of fixed alleles increased steadily over time, but the rate of fixation was faster for same than opposite alleles. Interestingly, fixation of opposite alleles was faster under high dominance, while fixation of same alleles was faster under low dominance degree. The divergence between the pools is primarily driven by the fixation of opposite alleles, which contributed to greater differentiation between the heterotic pools under higher dominance degree.

**Fig. 4.**
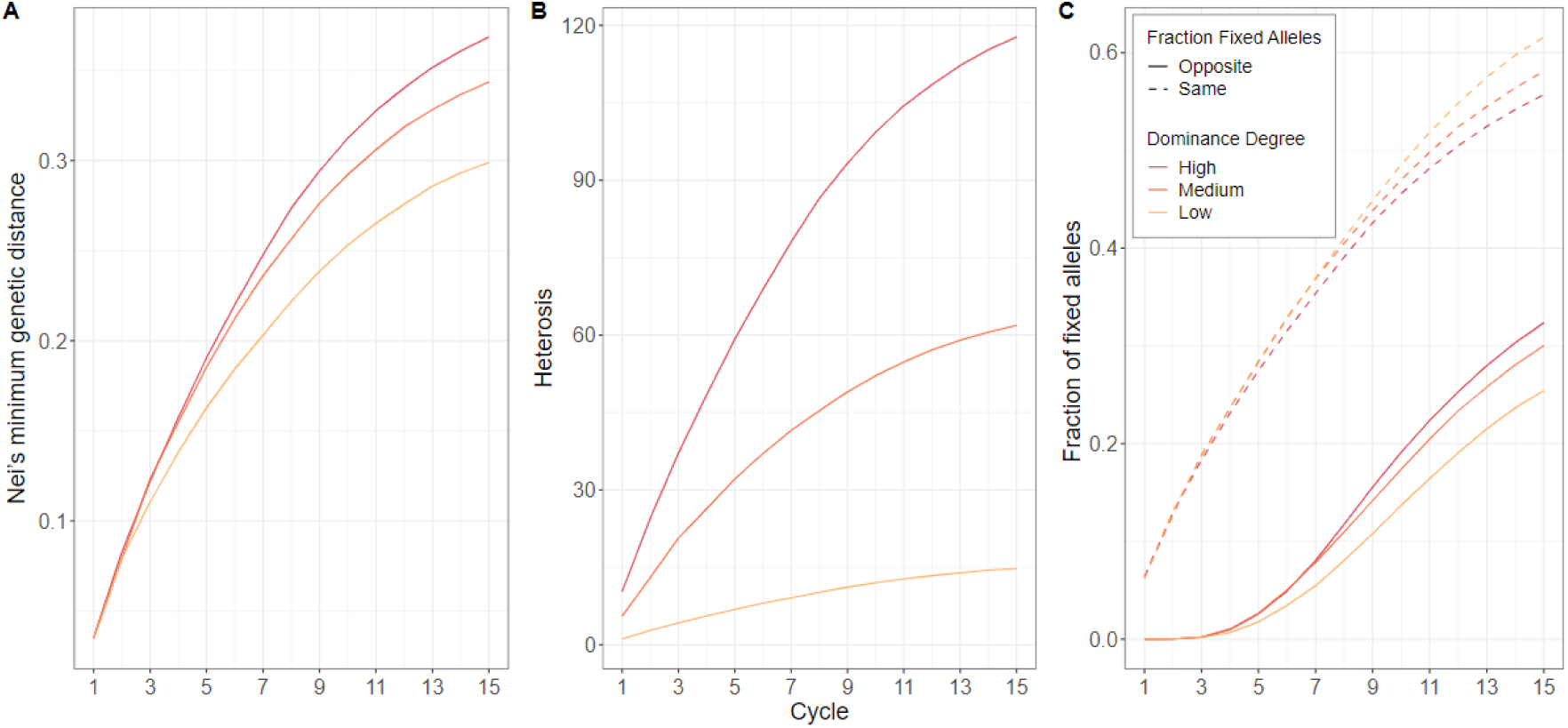
Metrics of genetic divergence between heterotic pools over 15 cycles of GCA-based selection in the baseline recurrent reciprocal genomic selection breeding program for high, medium and low degrees of dominance. A) Mean Nei’s minimum genetic distance, B) mean heterosis, and C) mean fraction of fixed (same or opposite) alleles in the two pools. The results are averaged across all 100 simulation replications of the baseline breeding program

**Fig. 5.**
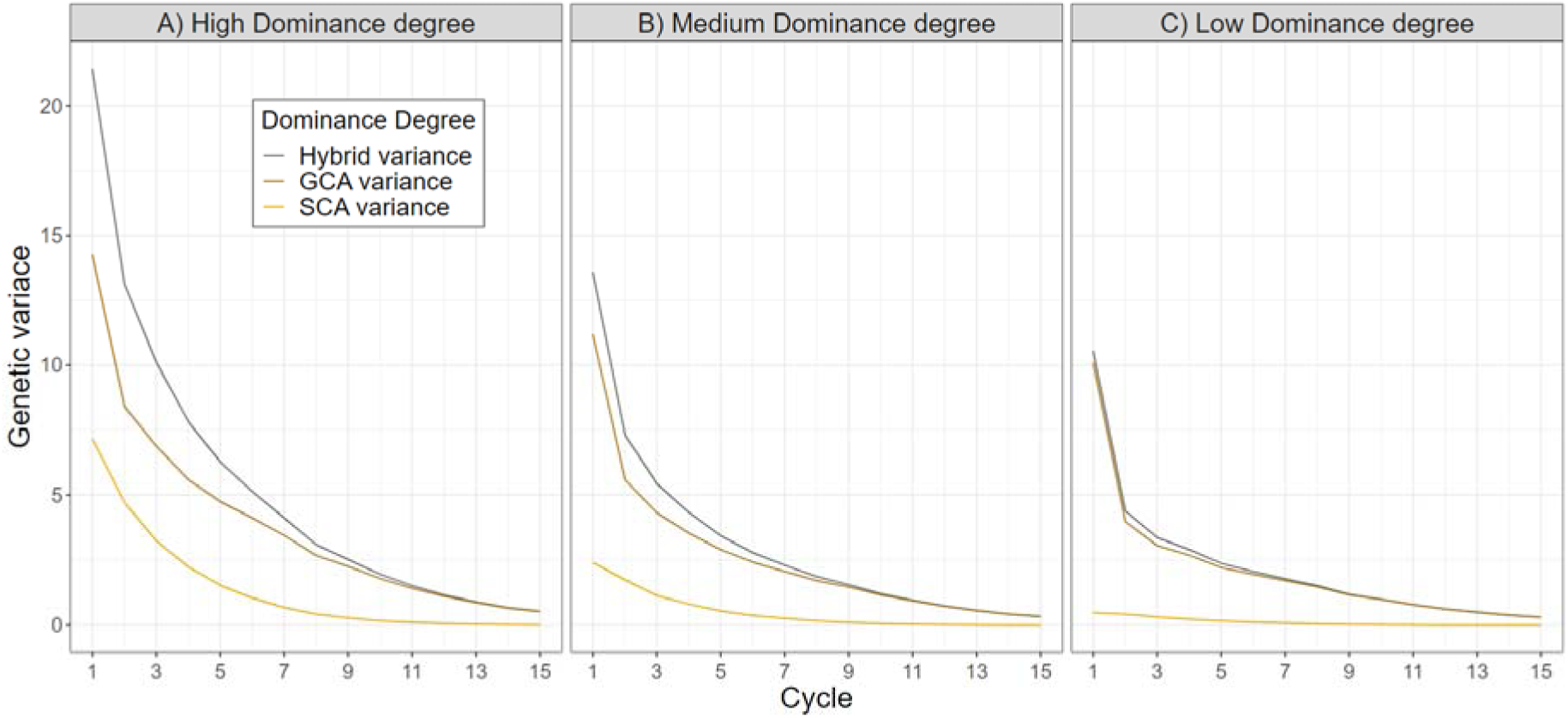
Hybrid genetic variance, GCA variance, and SCA variance monitored across 15 cycles of selection in the baseline recurrent reciprocal genomic selection breeding program, for A) high, B) medium and D) low degrees of dominance. The results are averaged across all 100 simulation replications of the baseline breeding program

Additionally, genetic diversity within pools decreased progressively as a consequence of selection. This was reflected by a faster increase in the Jaccard similarity coefficient over cycles (Suppl. Fig. S3B). Furthermore, the number of families used for within-pool crosses initially increased during the early selection cycles (2–5) before exhibiting a subsequent decline (Suppl. Fig. S3A). These results align with trends observed for genetic divergence, genetic variance components, and the fraction of fixed alleles.

### 4.2 Prediction accuracy of general combining ability

All 10 sparse testcrossing designs demonstrated higher genomic prediction accuracy for GCA compared to the conventional early-stage testcross design that utilizes a single tester. This advantage was most pronounced during the early breeding cycles but decreased as the genetic distance between pools increased. Figure 6 illustrates this, plotting the phenotypic and genomic GCA prediction accuracy at medium degree of dominance for the sparse testcrossing designs with sets of 2, 3, 4, 5, 10 and 20 testers, alongside the conventional early-stage testcross design with one tester, at cycles 1, 2, 3, 5, 10, and 15. Compared to phenotypic selection, genomic selection improved GCA accuracy by ∼0.3 (cycle 1) to ∼0.2 (cycle 15). Similar trends were observed at high (Suppl. Fig. S4) and low (Suppl. Fig. S6) dominance degrees.

**Fig. 6.**
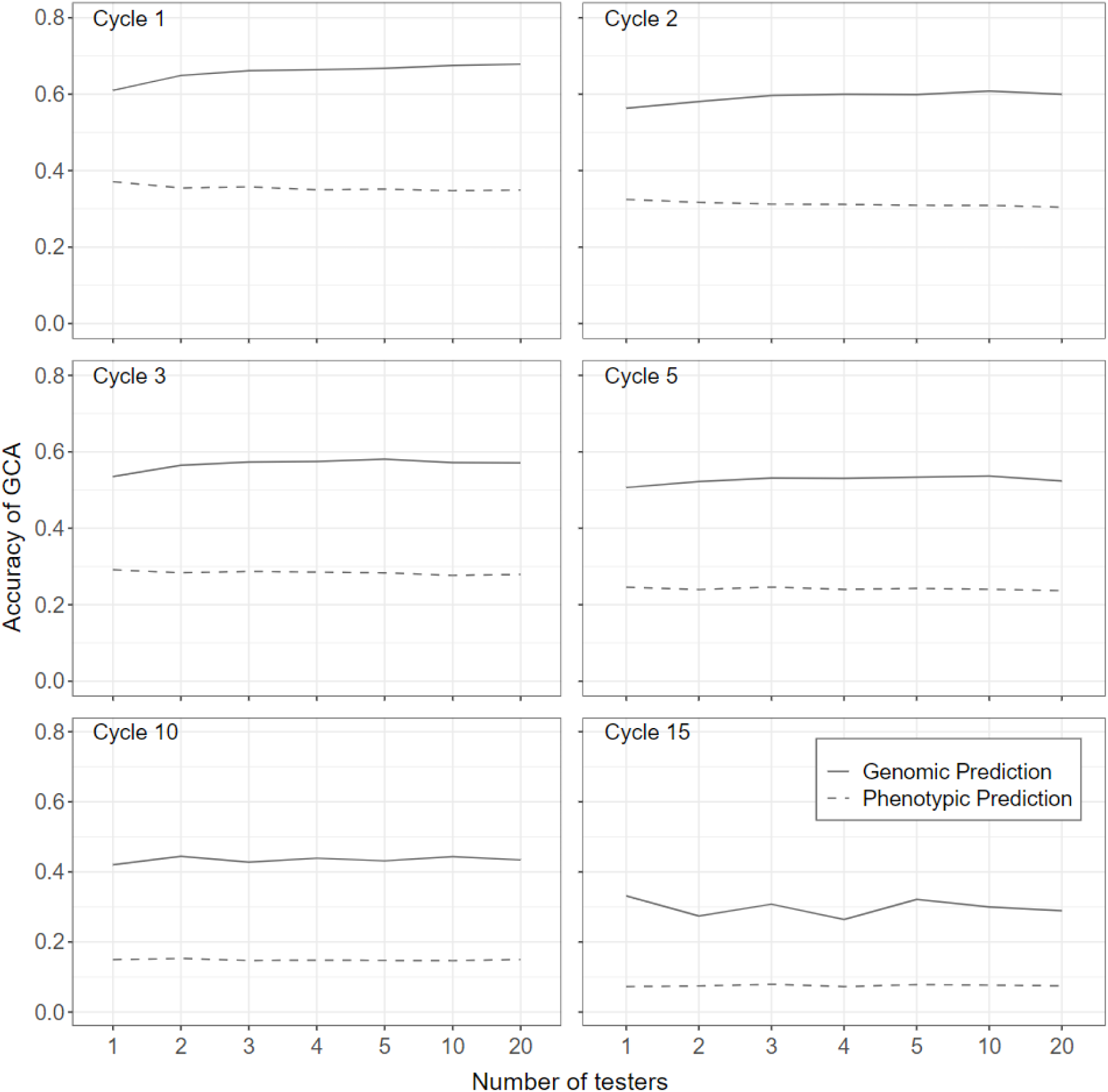
Phenotypic and genomic prediction accuracy of general combining ability (GCA) for early-stage candidate inbred lines using sparse testcrossing designs with sets of 2, 3, 4, 5, 10 and 20 testers, compared to a conventional early-stage testcross design with a single-tester. These designs were evaluated at selection cycles 1, 2, 3, 5, 10, and 15, and for a medium dominance degree within the female heterotic pool. The results are averaged across all 100 simulation replications of the baseline breeding program

Figure 6 also shows that in the early cycles, prediction accuracy improved with the number of testers, reaching a plateau around 5-10 testers, beyond which no significant increases were observed. For instance, when 10 testers were used, accuracy increased by approximately 3%, 11%, and 23% for low, medium, and high degrees of dominance, respectively, compared to the conventional strategy with one tester. Most of this accuracy increase was achieved with two or three testers. Since no further improvements were observed exceeding 10 testers, the results including sets of more than 20 testers for medium dominance degree were moved to Suppl. Fig. S5.

In contrast, by cycle 15, increasing the number of testers did not yield any further enhancement in prediction accuracy. These trends are further illustrated in Fig. 7, which plots the development of genomic GCA prediction accuracy for the conventional early-stage testcross design with one tester and the sparse testcrossing designs with sets of 2, 3, 4, 5, 10 and 20 testers against breeding cycles at high, medium and low degrees of dominance.

**Fig. 7.**
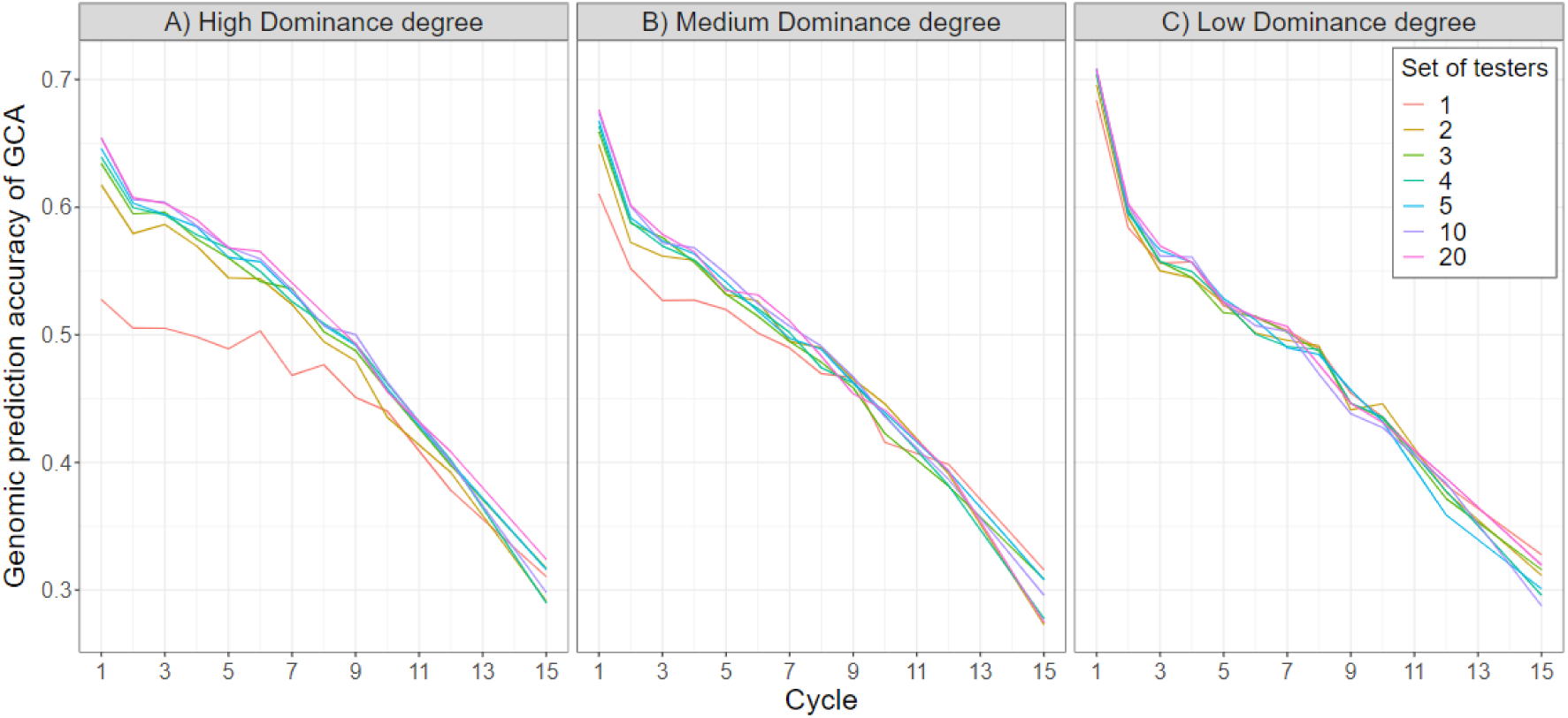
Genomic prediction accuracy of general combining ability (GCA) across 15 cycles of selection using sparse testcrossing designs with sets of 2, 3, 4, 5, 10 and 20 testers, compared to a conventional early-stage testcross design with a single-tester. Prediction accuracy was averaged across the female and male heterotic pools and for A) high, B) medium and C) low dominance degrees. The results are averaged across all 100 simulation replications of the baseline breeding program

Compared to Fig. 6, this figure focuses on dominance degrees and their effect on genomic prediction accuracy across cycles.

Figure 7 further indicates that genomic prediction accuracy improved more with sparse testcrossing at a high degree of dominance. However, when the dominance degree was low, there were no clear differences between the sparse testcrossing designs and the conventional early-stage testcross strategy that uses a single tester.

Additionally, Figures 6 and 7 reveal a decline in prediction accuracy over time for all sparse testcrossing designs as well as the conventional testcross design using a single tester. This decline can be attributed to a reduction in broad-sense heritability (Suppl. Fig. S7), which results from a decrease in hybrid genetic variance due to selection (Fig. 5), while the error variance remained constant over time. The phenotypic selection accuracy of GCA was consistently significantly lower than the genomic prediction accuracy of GCA and was not influenced by the testcross strategy (Fig. 6). Notably, phenotypic selection accuracy dropped to nearly zero in the final cycle across all degrees of dominance, while genomic prediction accuracies remained around (∼0.3), highlighting the reliability of genomic selection in later cycles.

### 4.3 Hybrid genetic gain

Hybrid genetic gain in the simulated recurrent reciprocal genomic selection breeding programs was consistently higher when sparse testcrossing was used to predict genomic GCA compared to the conventional early-stage testcross design with a single tester. This advantage was more pronounced at higher degrees of dominance but diminished as the degree of dominance decreased. Figure 8 illustrates this, by plotting hybrid genetic gain against breeding cycles for the conventional early-stage testcross design with one tester alongside the sparse testcrossing designs with sets of 2, 3, 4, 5, 10 and 20 testers. For example, with 10 testers, hybrid genetic gain in cycle 15 was approximately 3% higher than with one tester at low dominance degree, 6% higher at medium dominance degree, and 11% higher at high dominance degree.

**Fig. 8.**
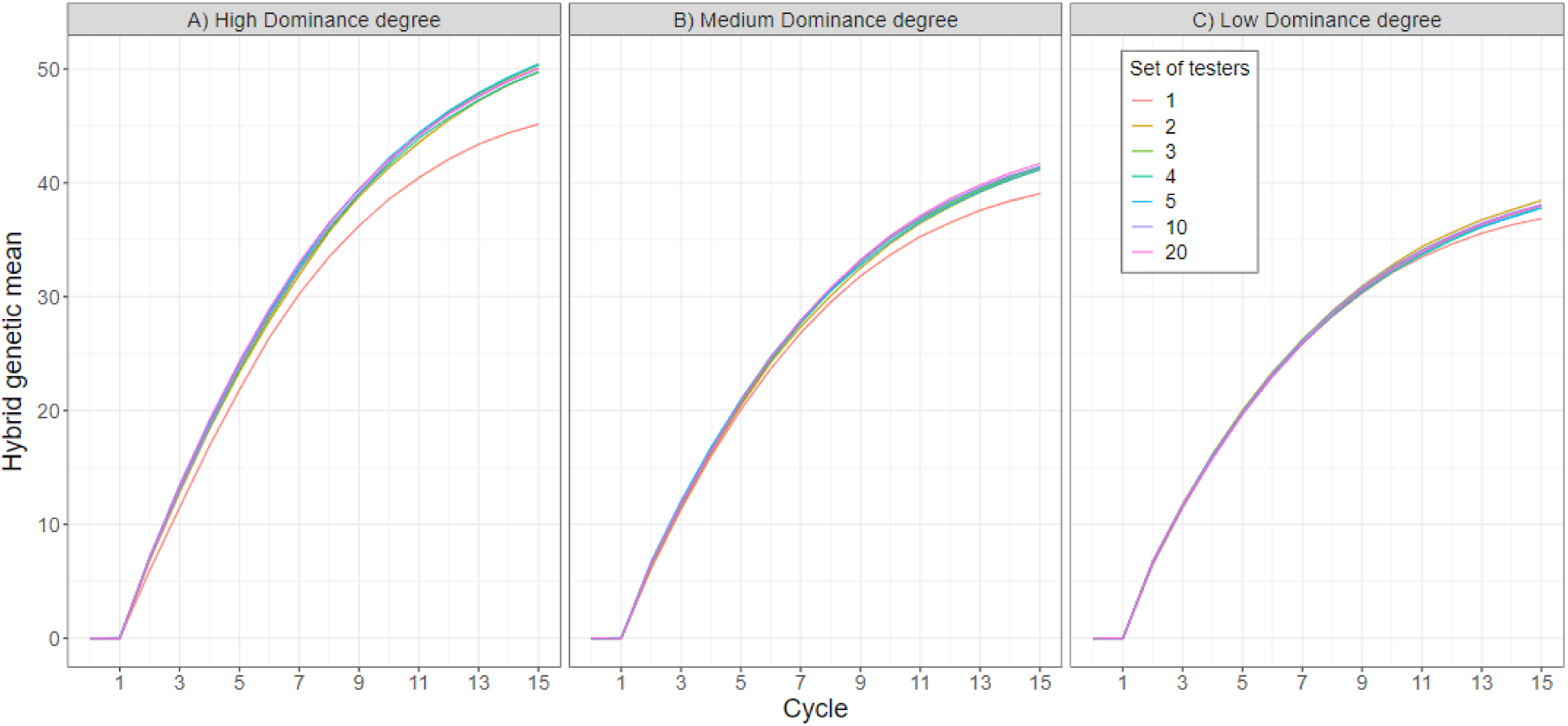
Accumulated hybrid genetic gain by using sparse testcross design with sets of 2, 3, 4, 5, 10 and 20 testers, compared to a conventional early-stage testcross design with a single-tester. This analysis was conducted across 15 cycles of recurrent reciprocal genomic selection and for A) high, B) medium and C) low dominance degrees. The results are averaged across all 100 simulation replications

Figure 8 also shows that hybrid genetic gain in cycle 15 improved with an increase in the number of testers. However, the differences in genetic gain among the different sparse testcrossing designs were relatively minor. For example, under medium dominance, genetic gain increased by 5.3% when using 2 testers compared to 1 tester, but only by 0.7% when comparing 10 testers to 2 testers. Furthermore, increasing the number of testers beyond 10 resulted in only marginal or no further increase in genetic gain. This finding aligns with the plateau in prediction accuracy around 5-10 testers observed in Fig. 6. Since no further improvements were observed exceeding 10 testers, the results for sets of more than 20 testers were moved to supplementary Fig. S8.

Interestingly, the sparse testcrossing designs and the conventional testcross design with a single tester showed similar increases in genetic distance and heterosis over time (Suppl. Fig S9 top and middle rows). The conventional testcross strategy, however, showed a faster increase in the fraction of fixed alleles in the two pools, in both same and opposite (Suppl. Fig S9 bottom row), and a slightly faster decline in GCA variance (Suppl. Fig S10). Furthermore, the conventional testcross strategy showed a lower within-pool genetic diversity as indicated by a faster increase in the Jaccard Similarity Coefficient (Suppl. Fig S11 top row). Additionally, the number of families within pools was consistently lower with a single tester compared to the sparse testcrossing designs and decreased slightly over time, while the number of families increased for the sparse testcrossing designs (Suppl. Fig S11 bottom row).

## 5 DISCUSSION

Sparse testcrossing using multiple testers can enhance genomic prediction accuracy of general combining ability (GCA) compared to conventional early-stage testcross strategies using a single tester, thus enabling increased rates of hybrid genetic gain. To discuss these results and their implications, the discussion section is divided into two parts. In the first part we explain that a relatively low number of three to five testers is sufficient for sparse testcrossing to be effective. We also emphasize that sparse testcrossing has the most significant impact on GCA prediction accuracy and hybrid genetic gain during the early stage of heterotic pool formation, particularly for traits where dominance is important. In the second part, we present how sparse testcrossing offers a promising alternative to conventional testcross designs and incomplete factorial designs.

### 5.1 Sparse Testcrossing Enhances GCA Prediction Accuracy and Hybrid Genetic Gain by employing Multiple Testers

All sparse testcrossing designs demonstrated higher genomic prediction accuracy for GCA compared to the conventional early-stage testcross strategy that utilizes a single tester, leading to increased hybrid genetic gain (Figs. 6 and 8). This improvement arose because sparse testcrossing, through the use of a genomic relationship matrix, enabled the evaluation of a genotype’s combining ability against multiple testers without increasing testcrossing resources. As a result, this method more effectively captured the genetic diversity of the heterotic pool than relying on a single tester.

In the early stages of hybrid breeding programs, a significant number of candidate lines are usually assessed for their GCA, which places considerable demands on testing resources. To efficiently use resources and maximize the number of candidate lines assessed, lines are often testcrossed with just one tester in the first stage of agronomic testing (Stage 1).

Only as the number of genotypes decreases in later stages, a conventional testcross design with multiple testers is used to enhance GCA prediction accuracy, as a larger pool of testers more accurately reflects the diversity of haploblocks and their frequencies within the complementary heterotic pool. Consequently, conventional hybrid breeding programs typically do not consider GCA to be reliably estimated after Stage 1 evaluation with a single tester, delaying the selection of candidates for recombination until after a second stage of evaluation with multiple testers. This approach, however, can extend the breeding cycle, potentially slowing genetic progress.

Sparse testcrossing enables the use of multiple testers in early-stage testing to predict the GCA of a genotype without increasing resource demands. This approach leverages genomic relationships, treating individual marker haplotypes as the primary unit of evaluation rather than the individual genotype. Although sparse testcrossing involves crossing each genotype with only a single tester, closely related genotypes with similar haplotypes can be crossed with different testers. This enables the evaluation of individual haplotypes, as well as common haplotype combinations, for their value and complementarity against a broader, more representative set of haplotypes from another heterotic pool carried by various testers. Consequently, the predicted values of the haplotypes in a single genotype reflect their average performance across multiple testcrosses rather than just one. This leads to higher GCA prediction accuracy, ultimately translating into greater genetic gain. Additionally, sparse testcrossing facilitates the selection of parents for recycling after the first stage, as it provides reliable GCA estimates. Recycling parents after Stage 1, rather than in later stages, can significantly shorten the breeding cycle. Thus, sparse testcrossing contributes to increased genetic gain by both improving the accuracy of GCA-based selection and reducing the duration of the breeding cycle.

In the absence of a genomic relationship matrix (GRM) to leverage information from relatives, phenotypic selection exhibited substantially lower prediction accuracy compared to the genomic prediction scenarios (Fig. 6). As result, sparse testcrossing without use of the GRM did not enhance prediction accuracy compared to using a single tester, since no information from relatives crossed with other testers contributed to a genotype’s predicted GCA. Additionally, testing lines with different testers can result in non-comparable GCA predictions and unfair comparisons. Therefore, it is not advisable to conduct sparse testcrossing without a GRM.

### 5.2 Sparse Testcrossing Enables enhanced GCA Prediction Accuracy with Few Testers

In the early selection cycles, we observed that the genomic prediction accuracy of GCA and hybrid genetic gain increased with the number of testers, reaching a plateau at around 5-10 testers (Figs. 6 and 8). As shown in Fig. 6, there is a minimal increase in prediction accuracy beyond three testers, regardless of the number of cycles. This suggests that using 3-5 testers for sparse testcrossing can effectively capture most of the diversity of haplotypes and haplotype combinations produced when crossed with another heterotic pool.

Surprisingly, increasing the number of testers beyond 10 did not result in any further appreciable gains in prediction accuracy. This may be because just a few cycles of crossing and selection in a closed breeding population can create strong linkage disequilibrium (Werner et al. 2023) and reduce within-pool haplotype diversity due to increased relatedness. As a result, a relatively small number of testers may sufficiently capture the most important haplotypes and haplotype combinations necessary for predicting GCA. Additionally, there may be a trade-off between testing a greater variety of haplotypes and haplotype combinations with the testers’ heterotic pool, and the replication rate of these combinations. While increasing the number of testers can enhance the representation of haplotype diversity, individual haplotypes may be tested less frequently, which leads to reduced accuracy in estimating their effects – especially for those that are in low frequency in the population and not well represented in many testcross combinations.

### 5.3 Sparse Testcrossing Enhances GCA Prediction Accuracy When Genetic Diversity Within Heterotic Pools Is High

Sparse testcrossing improved prediction accuracy and genetic gain most during the early breeding cycles. However, these advantages diminished as the genetic distance between pools increased over time (Figs. 6 and 7) and genetic diversity within pools decreased, as evidenced by the faster increase in the Jaccard similarity coefficient (Suppl. Fig. S3B).

As anticipated, the use of sparse testcrossing was most effective when there was high genetic diversity within heterotic pools, allowing for the capture of this diversity through multiple testers. This strategy resulted in a notable increase in prediction accuracy for GCA compared to the conventional testcross strategy with a single tester. However, as diversity within both pools decreased due to multiple cycles of selection, as indicated by an increase in the Jaccard coefficient (Suppl. Fig. S3B) and an increase in the fraction of fixed alleles (Fig. 4C), one or very few testers became sufficient to represent the reduced diversity of haplotypes and haplotype combinations in the testcrosses.

### 5.4 Sparse Testcrossing Enhances GCA Prediction Accuracy and Hybrid Genetic Gain When Dominance Is High

The improvement in GCA accuracy and genetic gain with sparse testcrossing was more pronounced when the degree of dominance was high but diminished as the degree of dominance decreased. The improvement is primarily due to the contribution of biological dominance effects to the general combining ability. The GCA captures both additive (*a*) and, to some extent, dominance genetic ( ) effect, as shown in the average testcross effects of an allele substitution (Eq. 3). Additionally, the extent to which GCA captures dominance depends on the magnitude of the dominance effects and the allele or haploblock frequencies in the tester pool. When dominance is high, dominance effects significantly contribute to GCA, with their impact depending on the allele frequencies within the tester pool. Therefore, sparse testcrossing using multiples testers improves the estimation of additive and dominance effects and the representation of allele or haploblock diversity and their frequencies of the tester pool, making a better exploitation of beneficial interaction between complementary alleles, thereby enhancing GCA accuracy. In contrast, when dominance is low, most part of GCA is attributable to additive effects, and the contribution from interaction between complementary alleles or haploblocks due to dominance becomes negligible. Consequently, increasing the number of testers does not substantially improve GCA accuracy.

### 5.5 Sparse Testcrossing Preserves Genetic Diversity Within Pools by Using Multiple Testers, thereby Enhancing Long-Term Hybrid Genetic Gain

The conventional early-stage testcross strategy using a single tester reached a faster plateau in hybrid genetic gain than the sparse testcrossing designs. This is primarily due to a more rapid decay of the GCA variance (Suppl. Fig. S10). Candidate lines that exhibit high GCA when crossed with the same single tester often share a common genetic background, benefiting from similar haploblock combinations and their interactions. Consequently, relying on a single tester increases the likelihood of selecting closely related lines, such as from the same family, unless this is actively managed using an approach such as optimal contribution selection (Woolliams et al. 2015).

In the conventional two-stages selection approach, although lines are selected as parents in Stage 2 by using multiple testers for more accurate prediction of GCA, most of the selection pressure is applied at Stage 1, where thousands of candidate lines are discarded based solely on data from a single tester. The use of a single tester imposes a severe bottleneck on early-stage genetic variation, reducing the genetic diversity of the subset advanced to Stage 2. This leads to a rapid accumulation of a somewhat restricted set of haplotypes within pools, which accelerates the decay of genetic variance and ultimately limits further genetic gain. Evidence for this is found in the faster increase of the Jaccard similarity coefficient (Suppl. Fig. S11 -top row-) and the high fraction of fixed alleles in the two pools (Suppl. Fig. S9 -bottom row-) in the conventional testcross strategy with a single tester. Additionally, the number of families used for crossing was consistently lower with a single tester and further decreased over selection cycles (Suppl. Fig. S11 -bottom row-). Particularly when genomic selection is applied, the combination of increased accuracy and rapid parent recycling in a closed breeding program can exacerbate the erosion of genetic diversity within pools.

Conversely, sparse testcrossing prioritizes lines with high GCA predicted from a more diverse set of testers at early-stage. This strategy effectively redirects selection pressure away from closely related genotypes that may perform well with the same single tester, toward a genetically more diverse set of candidates. As a result, sparse testcrossing not only enhances short-term genetic gains but also promotes a more sustainable and resilient breeding program, facilitating long-term genetic improvements without rapidly depleting genetic variance.

### 5.6 Sparse Testcrossing offers an Effective Balance between increased GCA Accuracy and Resource Efficiency compared to Incomplete Factorial Designs

Sparse testcrossing provides breeders with an effective balance between simple testcross designs, resource efficiency, and increased prediction accuracy for general combining ability (GCA), thereby laying the foundation for increased rates of genetic gain. It offers a promising alternative to conventional testcrossing strategies, where each candidate is crossed with the same tester(s), as well as incomplete factorial designs, which randomly sample testcrosses from a complete factorial mating design.

To compare our sparse testcrossing designs with an incomplete factorial testcrossing strategy, we simulated a scenario with 1200 testers, thereby approximating an incomplete factorial design. The GCA prediction accuracies with 1200 testers were comparable to those achieved using sparse testcrossing designs with 3–5 testers (Suppl. Figs S4, S5 and S6 for high, medium and low degree of dominance). Incomplete factorial designs use a large number of testers, based on the assumption that this approach provides a more accurate representation of the testers’ heterotic pool to increase GCA prediction accuracy. However, our simulations demonstrate that sparse testcrossing designs with few testers can effectively provide a good representation of the testers’ heterotic pool, increasing GCA accuracy at early-stage testing. Additionally, our results show that as genetic distance between pools increases and within-pool diversity decreases, fewer testers are needed to represent the reduced diversity.

Although incomplete factorial designs can offer the advantage of evaluating twice as many lines for a given number of testcrosses or requiring half the phenotyping effort for the same number of candidates (Lorenzi et al. 2022), their implementation poses significant logistical challenges. These arise primarily due to labor-intensive and costly manual pollination required to achieve the intended hybrid combinations (Seye et al. 2020; Lorenzi et al. 2022) and are further intensified by the need to synchronize flowering times and produce sufficient seed quantities for multi-environment evaluations. In contrast, sparse testcrossing with 3–5 testers provides GCA accuracies comparable to incomplete factorial designs and can be implemented through modified open-pollination setups or hand pollination, which is much less complex than the approach used in an incomplete factorial design.

To effectively implement sparse testcrossing designs, we recommend crossing full-sibs with the various testers in a balanced manner. This approach ensures that haplotypes shared among full-sibs are evaluated alongside a diverse range of haplotypes representative of the testers’ heterotic pool, facilitating a more thorough assessment of a genotype’s complementarity with other heterotic pools.

### **5.7** Limitations of this study

In this study, we simulated the population improvement component of a rapid cycle genomic selection hybrid breeding program using inbred lines as testers. However, public hybrid breeding programs may use intra-pool single or double crosses as testers instead, which combine the haploblocks of multiple inbred lines and exhibit a certain level of heterozygosity. We hypothesize that these testers could, per se, provide a better representation of the genetic diversity within a heterotic pool, potentially requiring a lower number of testers to achieve most of the accuracy. However, confirming this will require further investigation. A notable disadvantage of using single and double cross hybrids is that their generation takes more time than that of doubled haploid or inbred line testers (Longin et al. 2007), which will imply using testers from previous cycles, at the cost of delaying the updating of haplotype frequencies.

Furthermore, all genes (QTN) simulated in AlphaSimR are biallelic, and heterotic pools contain the same genes with identical effects, except for those few undergoing mutation. Additionally, we did not simulate epistatic effects due to the complexity and difficulty of incorporating a realistic representation of different epistatic interactions. As a result, our simulation approach may underestimate the genetic diversity and genotype-specific background effects within and between heterotic pools. In this context, a slightly higher number of testers may indeed provide better predictions of GCA, and the differences in accuracy and gain between, for example, 2 testers and 10 testers may be more pronounced than in our simulations. With these considerations in mind, we cautiously recommend using 3–5 tester lines, balancing practical implementability, design simplicity, and selection accuracy.

Lastly, in our simulations, we selected testers randomly at each cycle to avoid confounding our results with the effects of specific tester selection methods. Across the 100 simulation replications, this ensured that different scenarios had equal chances of randomly drawing more or less representative sets of testers, providing on average a robust estimate of accuracy and gain. In conventional hybrid breeding programs, a more targeted approach to tester selection is typically employed. Practically speaking, testers are usually lines currently used in hybrids. They are often chosen based on high GCA values, superior *per se* performance, and desirable pollen characteristics, as these traits are critical for their role as parents of commercial hybrids (Melchinger and Frisch 2023). In some public programs, such as the CIMMYT maize breeding programs in Eastern and Southern Africa, testers are also selected for seed production characteristics when serving as single-cross female parents of three-way cross hybrids. This approach may not adequately capture the haplotype diversity of a heterotic pool. In fact, theoretical and experimental studies suggest that using low-performing testers with low frequencies of favorable alleles could be more effective for improving GCA (Rawlings and Thompson 1962; Hallauer et al. 2010). We recognize that the selection and updating of testers is a significant research question in its own right and is beyond the scope of this paper.

## 6 ACKNOWLEDGMENTS

## 7 SUPPLEMENTARY INFORMATION

Supplemental figures S1 to S12 are available in the Supplementary Material.

## 8 DECLARATIONS

### Author Contribution Statement

All authors contributed to the study conception. DOGD and CRW designed the simulation framework. DOGD scripted the simulations, did the analysis and generated the figures. DOGD and CRW wrote the first draft of the manuscript. All authors critically revised and contributed on previous versions of the manuscript. All authors read and approved the final manuscript.

## Data Availability

Not applicable

### Ethics approval

Not Applicable.

### Consent to participate

Not applicable.

### Consent to publication

Not applicable.

### Human and animal rights

Not applicable.

### Code availability

All simulations were done using AlphaSimR version 1.4.2 (Gaynor et al., 2021). The R-scripts used to generate the results presented in this simulation study are available on the GitHub repository at https://github.com/dogrdc/Sparse-testcrossing.

