## Supplemental Figures for "Sparse testcrossing for early-stage genomic prediction of general combining ability to increase genetic gain in maize hybrid breeding programs"

[1]Breeding Innovation and Modernization, Consultative Group of International Agricultural Research (CGIAR), Texcoco, Mexico.
[2] International Maize and Wheat Improvement Center (CIMMYT), Texcoco, Mexico.
[3] International Maize and Wheat Improvement Center (CIMMYT), Nairobi, Kenya.
[4]International Maize and Wheat Improvement Center (CIMMYT), Harare, Zimbabwe.
[5] Bill & Melinda Gates Foundation, Seattle, WA 98109, USA

**Corresponding author:**
David O. González-Diéguez
International Maize and Wheat Improvement Center (CIMMYT), Km 45, Carretera Mexico-Veracruz, Texcoco 56237, Edo. de México, Mexico


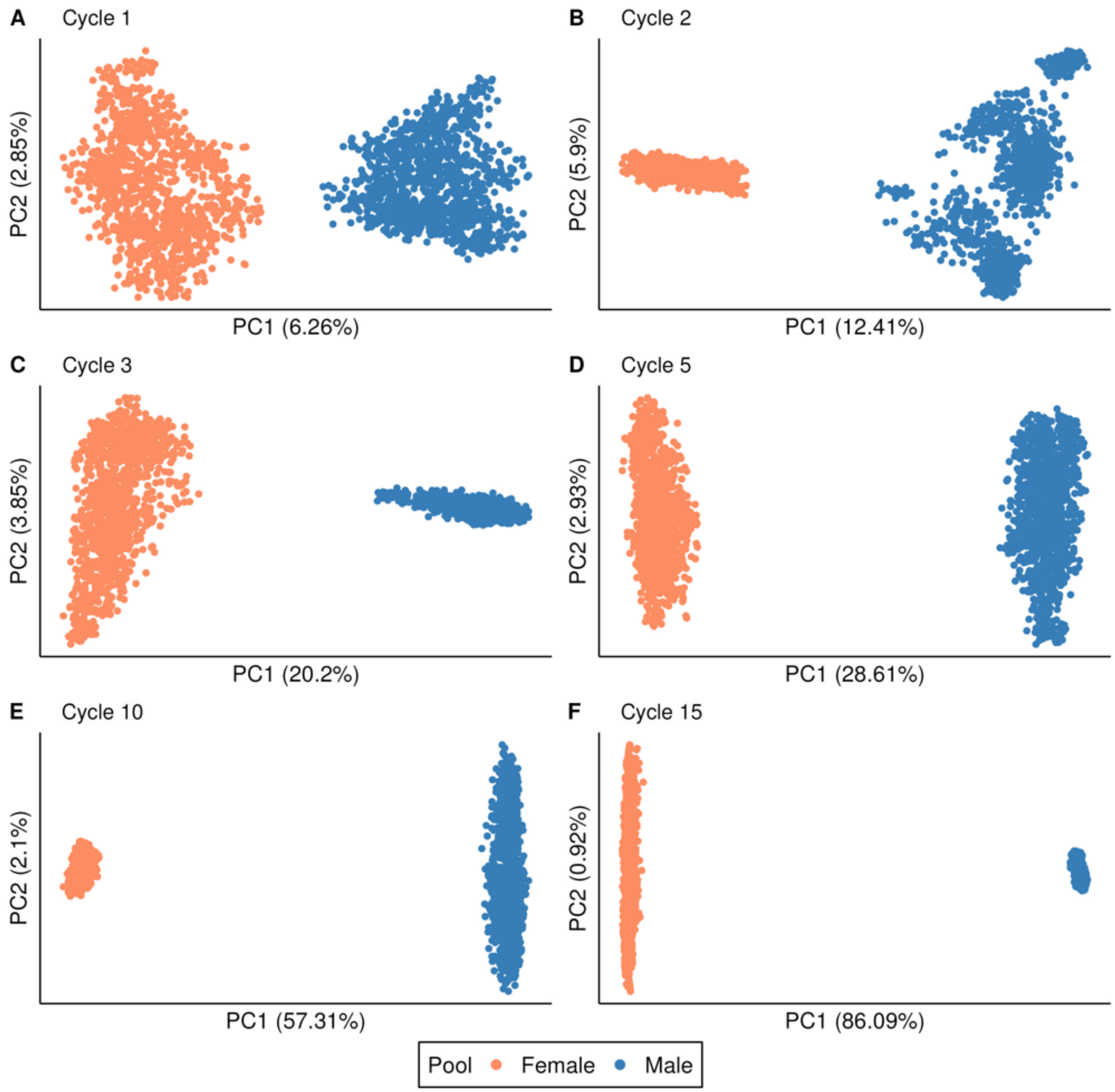


**Fig. S1** The first two principal components from a principal component analysis (PCA) performed on the genomic relationship matrix of the candidate inbred lines at cycles 1, 2, 3, 5, 10, and 15 of the baseline breeding program. The first principal component is represented on the X-axis, while the second principal component is plotted on the Y-axis. Colors orange and blue represent female and male heterotic pools, respectively. The proportion of variance explained by the first and second principal components (PC) is indicated in brackets. Results correspond to a single, randomly selected simulation replication of the baseline breeding program with low dominance degree

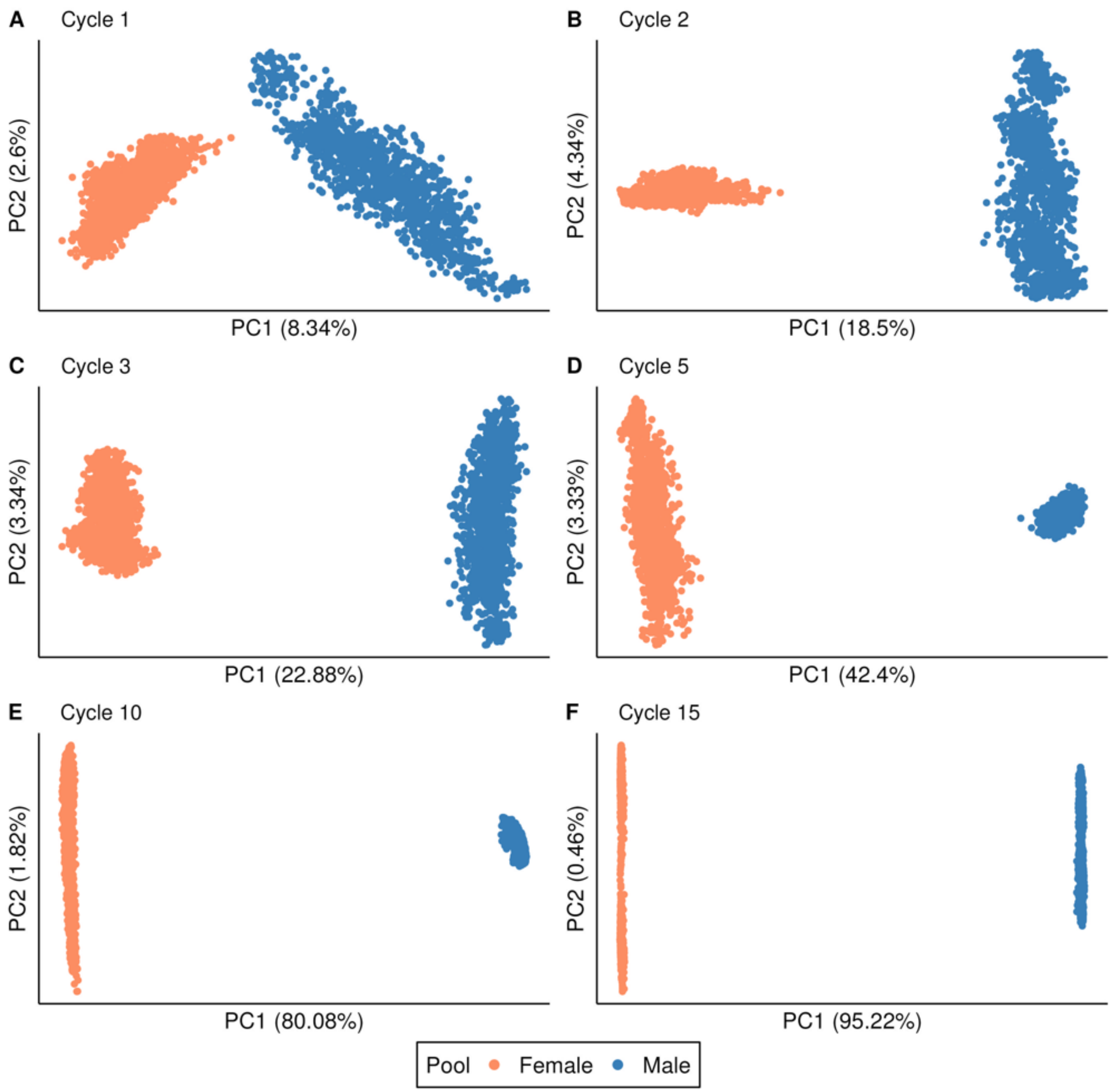


**Fig. S2** The first two principal components from a principal component analysis (PCA) performed on the genomic relationship matrix of the candidate inbred lines at cycles 1, 2, 3, 5, 10, and 15 of the baseline breeding program. The first principal component is represented on the X-axis, while the second principal component is plotted on the Y-axis. Colors orange and blue represent female and male heterotic pools, respectively. The proportion of variance explained by the first and second principal components (PC) is indicated in brackets. Results correspond to a single, randomly selected simulation replication of the baseline breeding program with high dominance degree

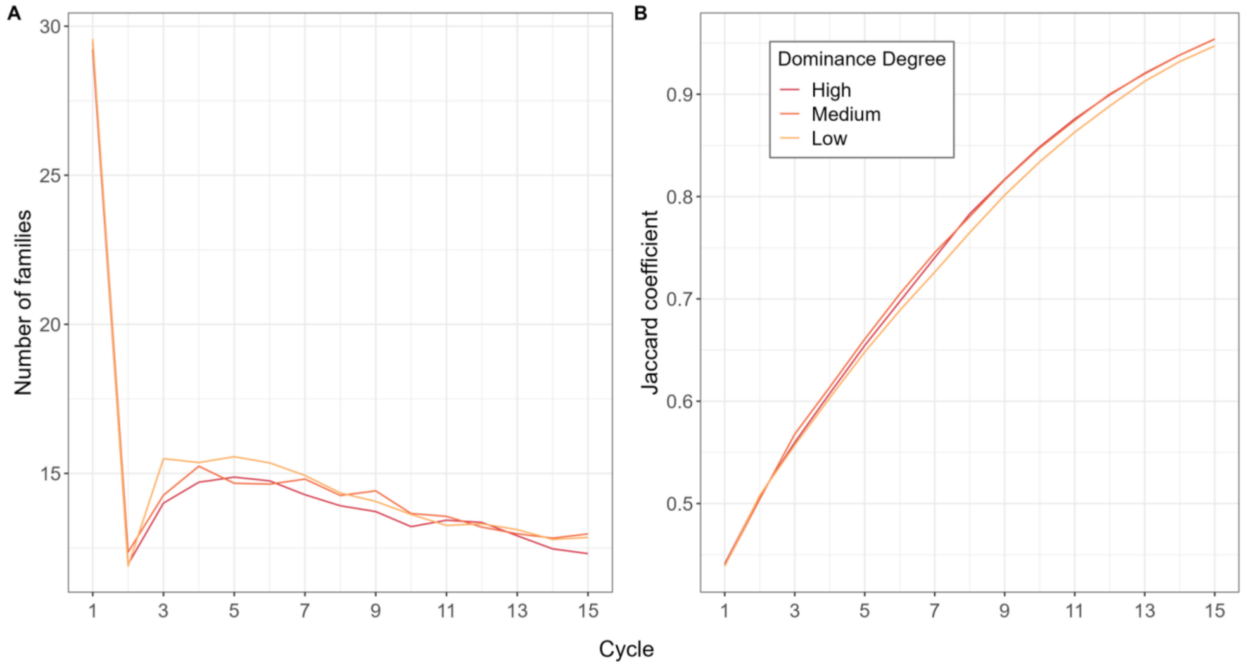


**Fig. S3** Metrics of genetic diversity within the breeding pools monitored across 15 cycles of selection in the baseline recurrent reciprocal genomic selection breeding program for high, medium and low dominance degrees. A) Number of families averaged across both female and male heterotic pools, B) Jaccard similarity coefficient averaged across pools. Results are averaged across all 100 simulation replications of the baseline breeding program

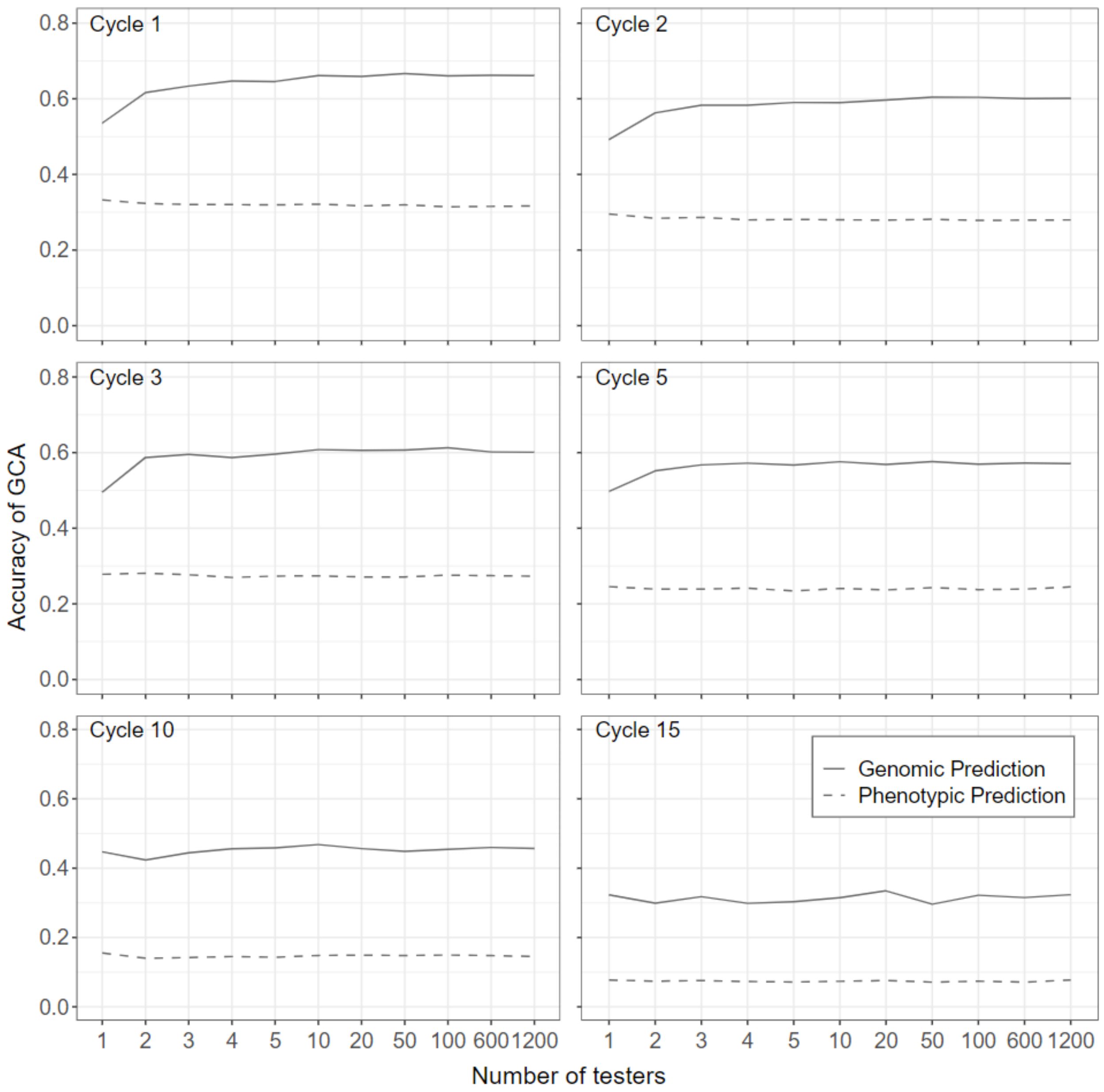


**Fig. S4** Phenotypic and genomic prediction accuracy of GCA effects for early-stage candidate inbred lines by using sparse testcross designs with sets of 2, 3, 4, 5, 10, 20, 50, 100, 600, and 1,200 testers, compared to a conventional early-stage testcross design with a single-tester, at selection cycles 1, 2, 3, 5, 10, and 15, and for a high dominance degree within the female heterotic pool. Results are averaged across all 100 simulation replications of the baseline breeding program

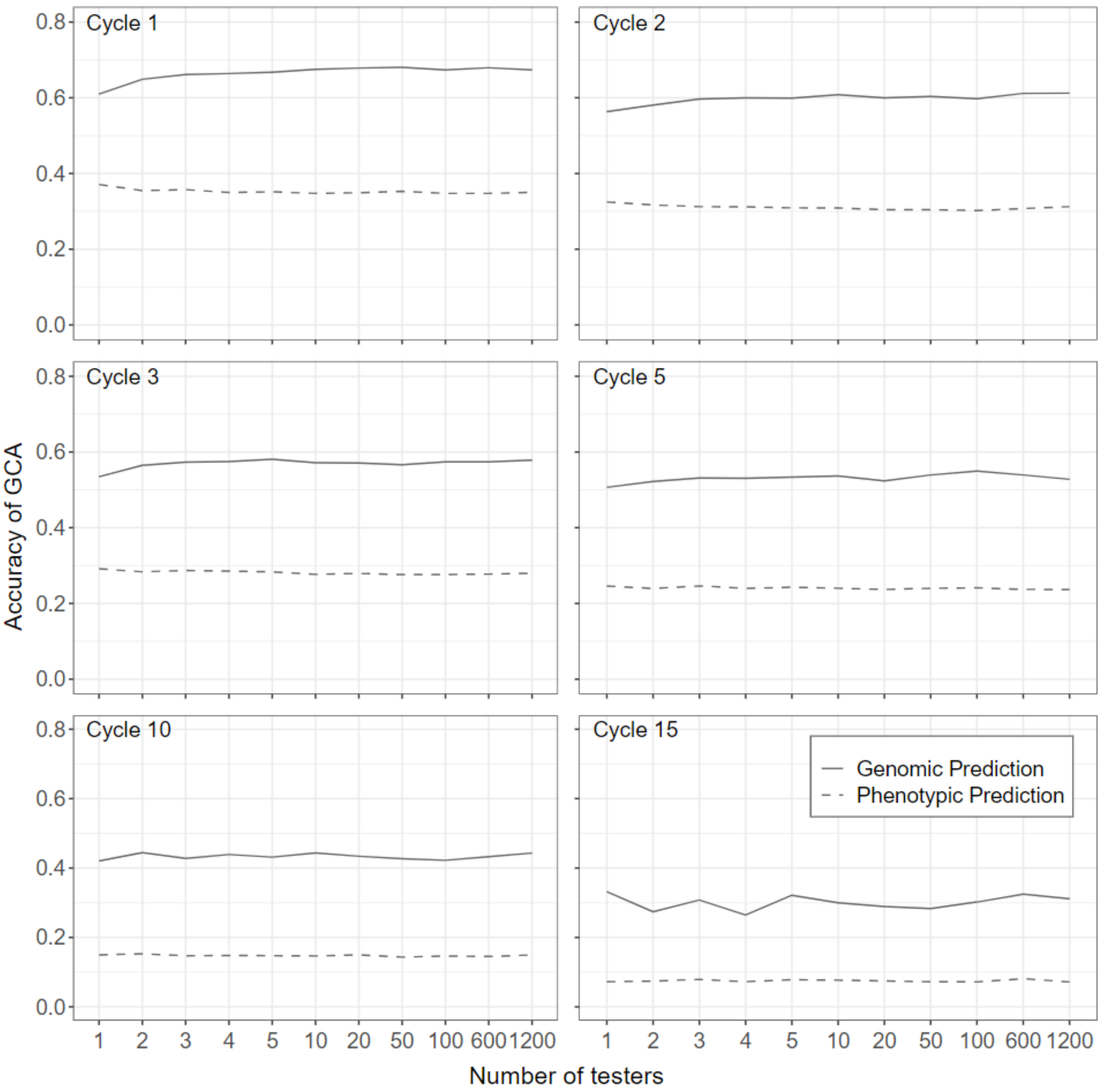


**Fig. S5** Phenotypic and genomic prediction accuracy of GCA effects for early-stage candidate inbred lines by using sparse testcross designs with sets of 2, 3, 4, 5, 10, 20, 50, 100, 600, and 1,200 testers, compared to a conventional early-stage testcross design with a single-tester, at selection cycles 1, 2, 3, 5, 10, and 15, and for a medium dominance degree within the female heterotic pool. Results are averaged across all 100 simulation replications of the baseline breeding program

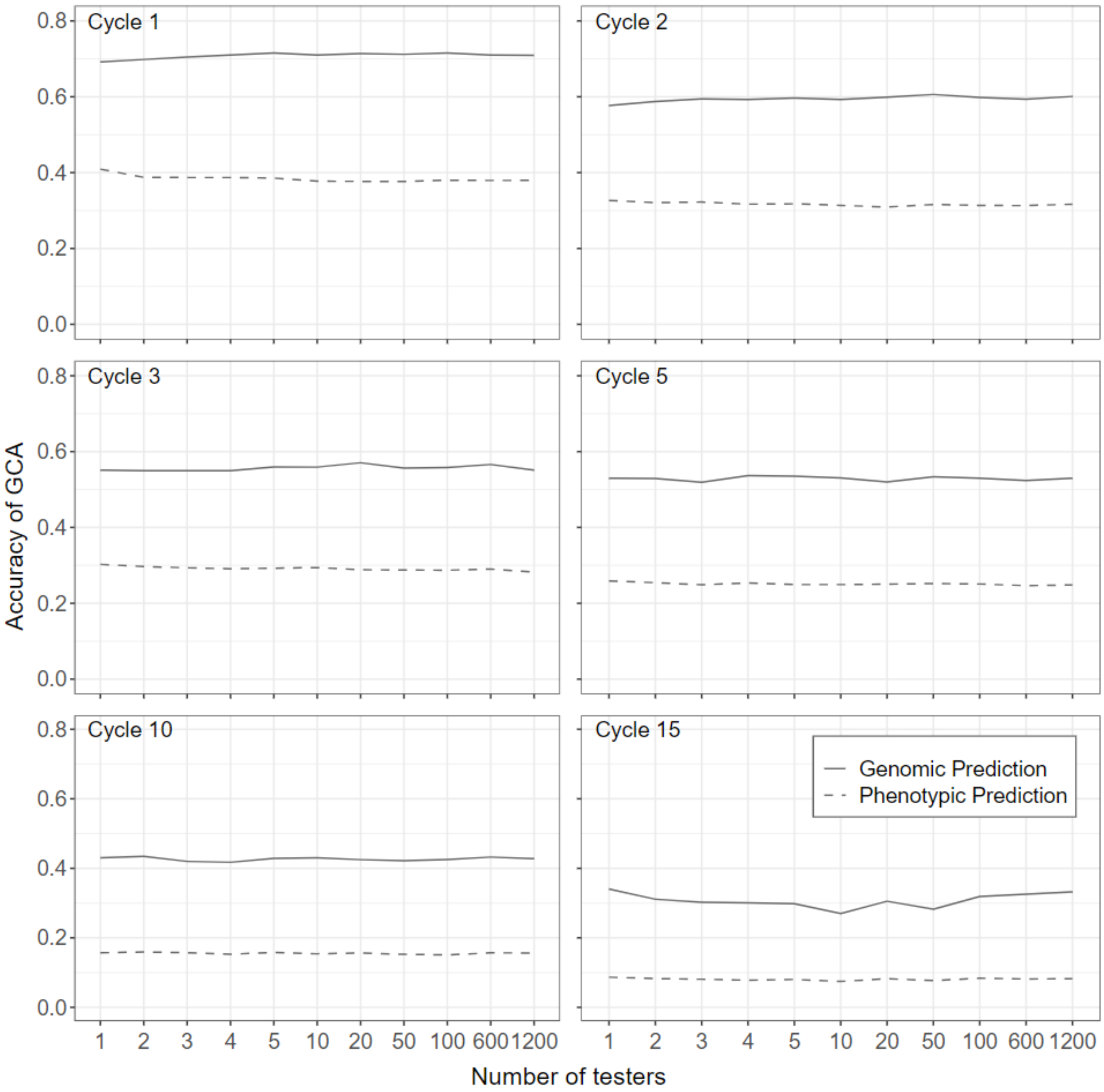


**Fig. S6** Phenotypic and genomic prediction accuracy of GCA effects for early-stage candidate inbred lines by using sparse testcross designs with sets of 2, 3, 4, 5, 10, 20, 50, 100, 600, and 1,200 testers, compared to a conventional early-stage testcross design with a single-tester, at selection cycles 1, 2, 3, 5, 10, and 15, and for a low dominance degree within the female heterotic pool. Results are averaged across all 100 simulation replications of the baseline breeding program

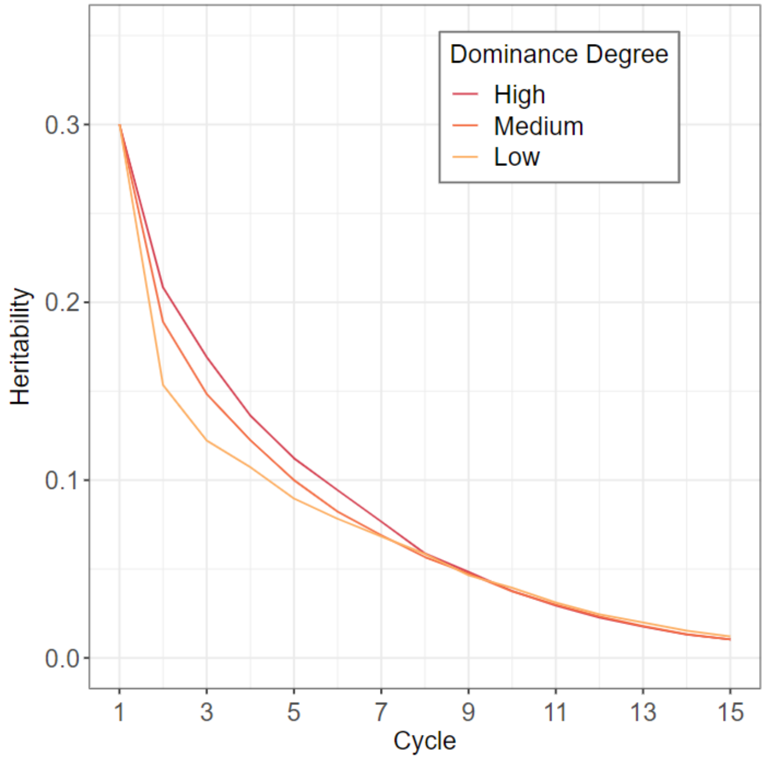


**Fig. S7** Broad-sense heritability ($H^2$) monitored across 15 cycles of selection in the baseline recurrent reciprocal genomic selection breeding program, for A) high, B) medium and D) low dominance degrees. Results are averaged across all 100 simulation replications of the baseline breeding program

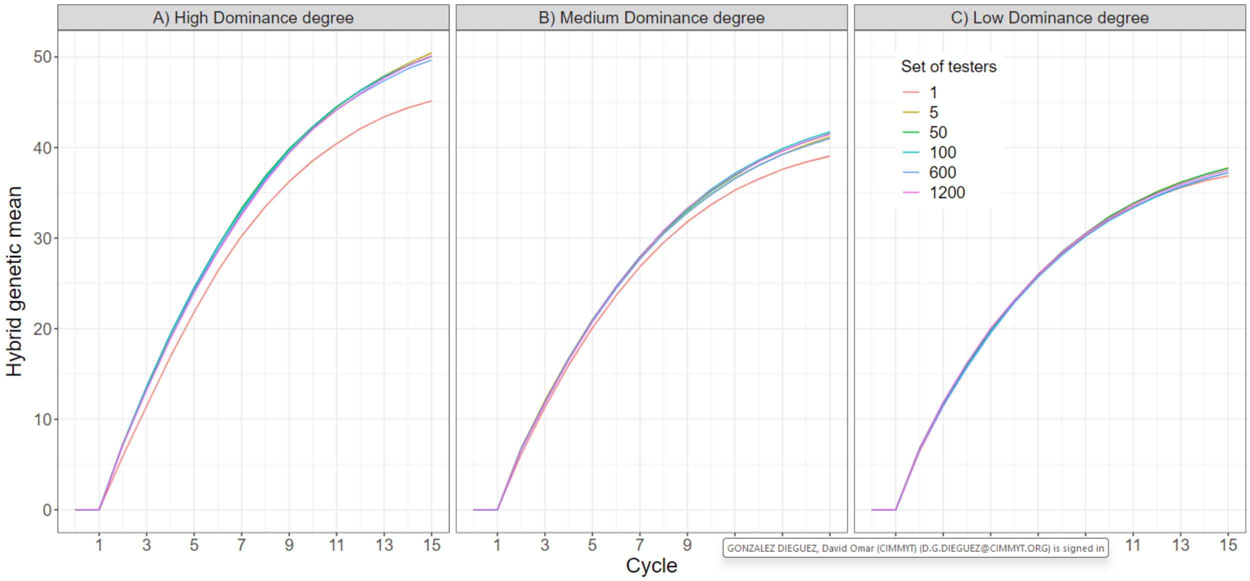


**Fig. S8** Accumulated hybrid genetic gain by using sparse testcross designs with sets of 5, 50, 100, 600 and 1200 testers, compared to a conventional early-stage testcross design with a single-tester, across 15 cycles of recurrent reciprocal genomic selection for A) high, B) medium and C) low dominance degree. Results are averaged across all 100 simulation replications

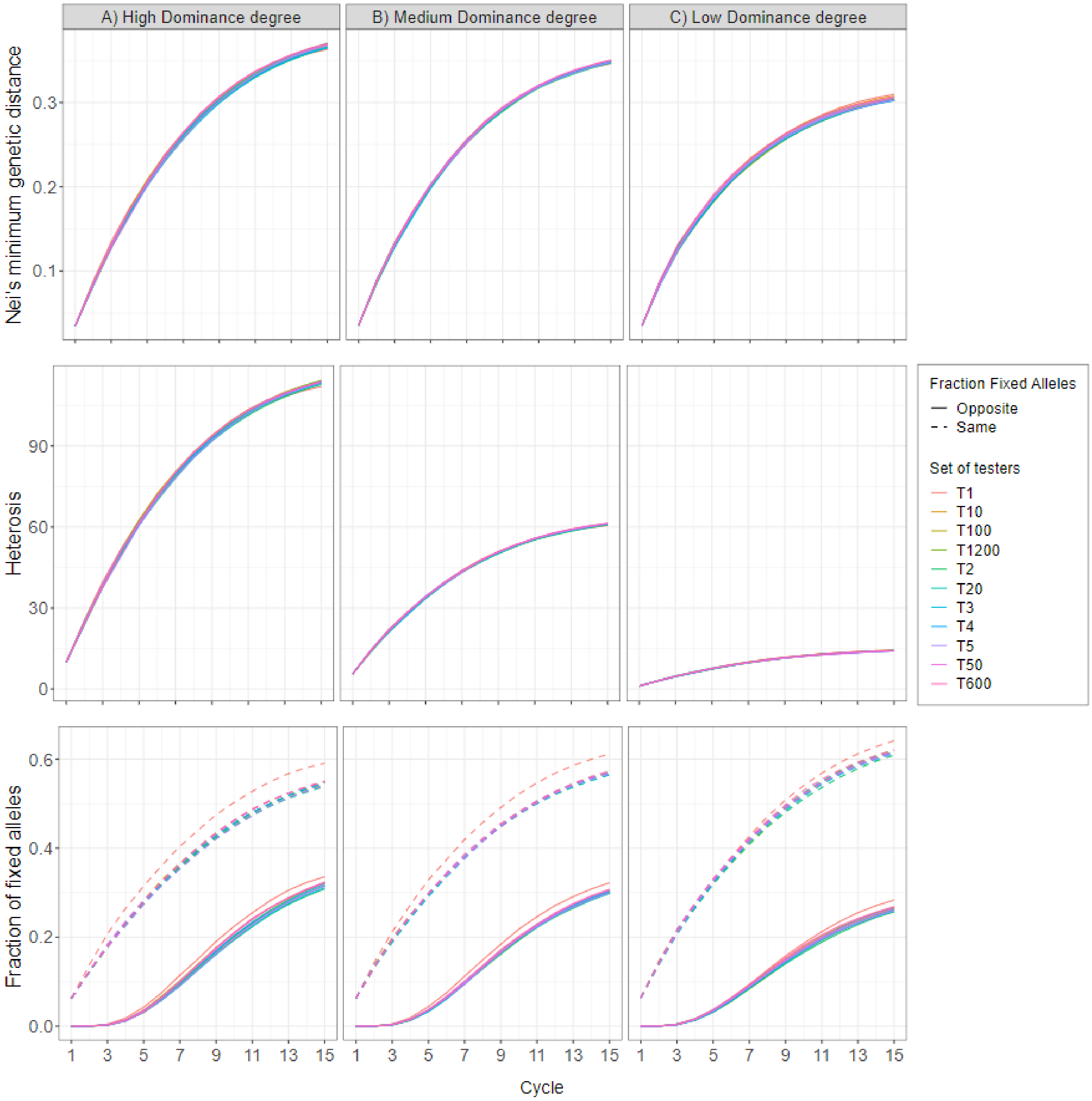


**Fig. S9** Metrics of genetic divergence between heterotic pools monitored across 15 cycles of selection for sparse testcross designs with sets of 2, 3, 4, 5, 10, 20, 50, 100, 600 and 1200 testers, and the conventional testcross design with a single-tester. Mean Nei's minimum genetic distance between heterotic pools, mean heterosis and mean fraction of fixed (same and opposite) alleles in the two heterotic pools, for A) high, B) medium and c) low dominance degrees. Results are averaged across all 100 simulation replications

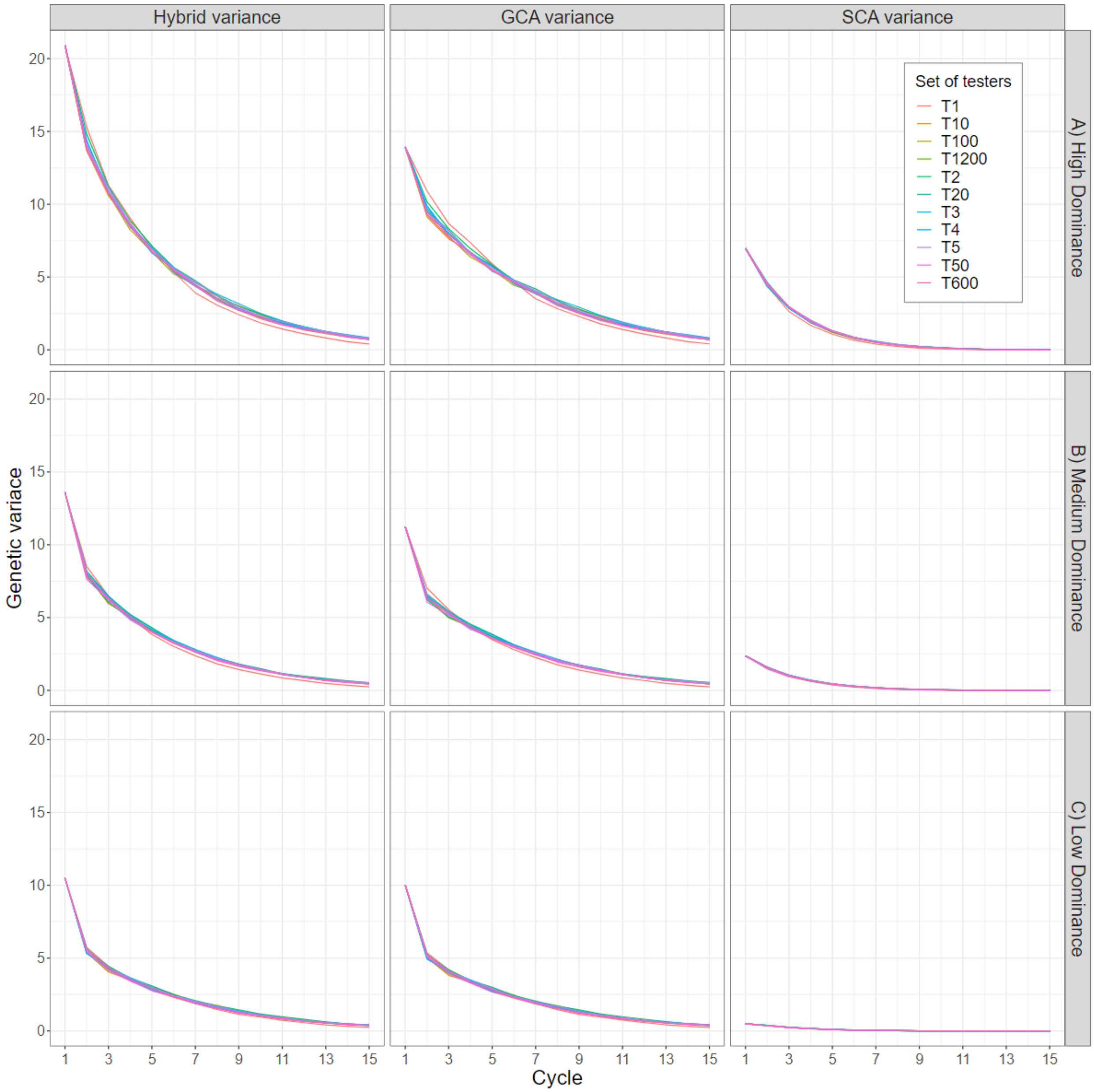


**Fig. S10** Total genetic variance of hybrids, GCA variance of inbred lines, and SCA variance in hybrids observed by using sparse testcross designs with sets of 2, 3, 4, 5, 10, 20, 50, 100, 600 and 1200 testers, compared to a conventional early-stage testcross design with a single-tester, across 15 cycles of recurrent reciprocal genomic selection and for A) high, B) medium and C) low dominance degrees. Results are averaged across all 100 simulation replications

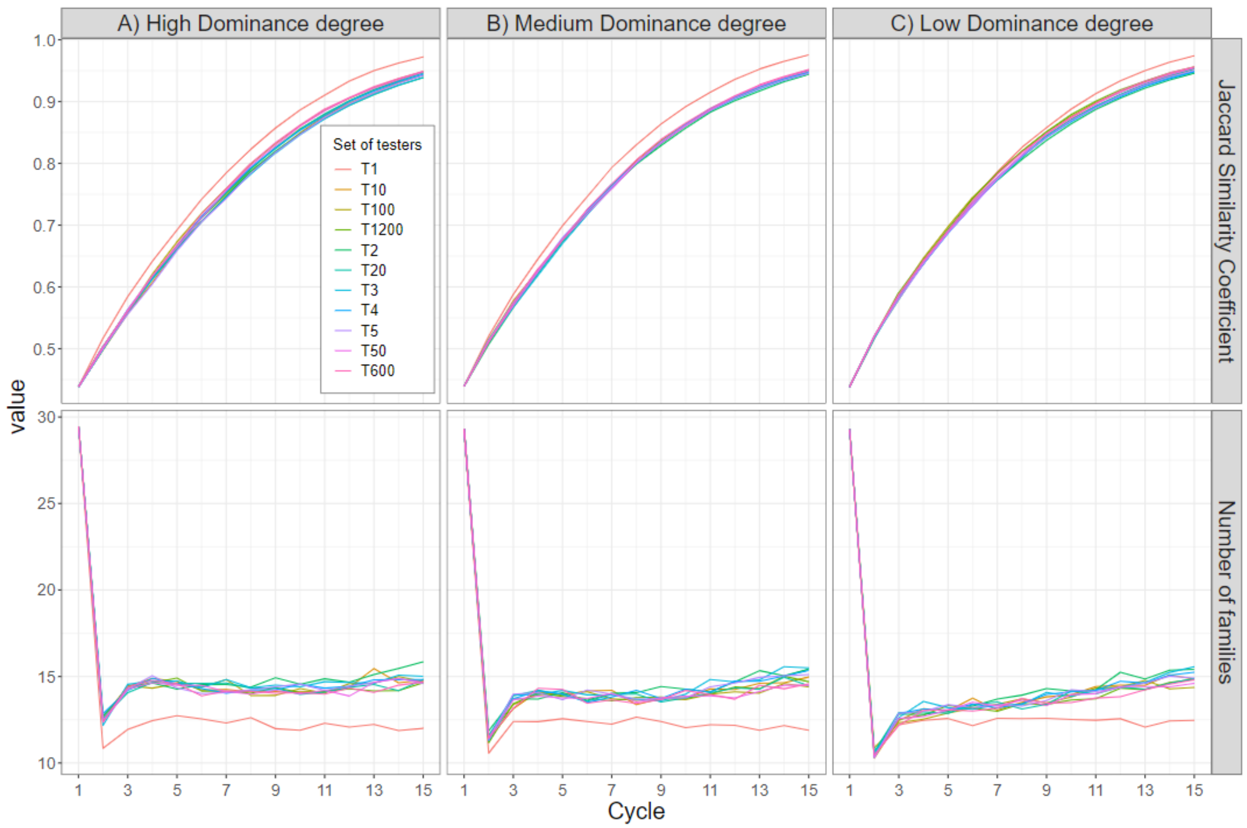


**Fig. S11** Metrics of genetic diversity within the breeding pools monitored across 15 cycles of selection for sparse testcross designs with sets of 2, 3, 4, 5, 10, 20, 50, 100, 600 and 1200 testers, and the conventional testcross design with a single-tester. Mean number of families used for crossing averaged across both female and male heterotic pools, and within pool Jaccard similarity coefficient averaged across male and female pools, for A) high, B) medium and c) low dominance degrees. Results are averaged across all 100 simulation replications

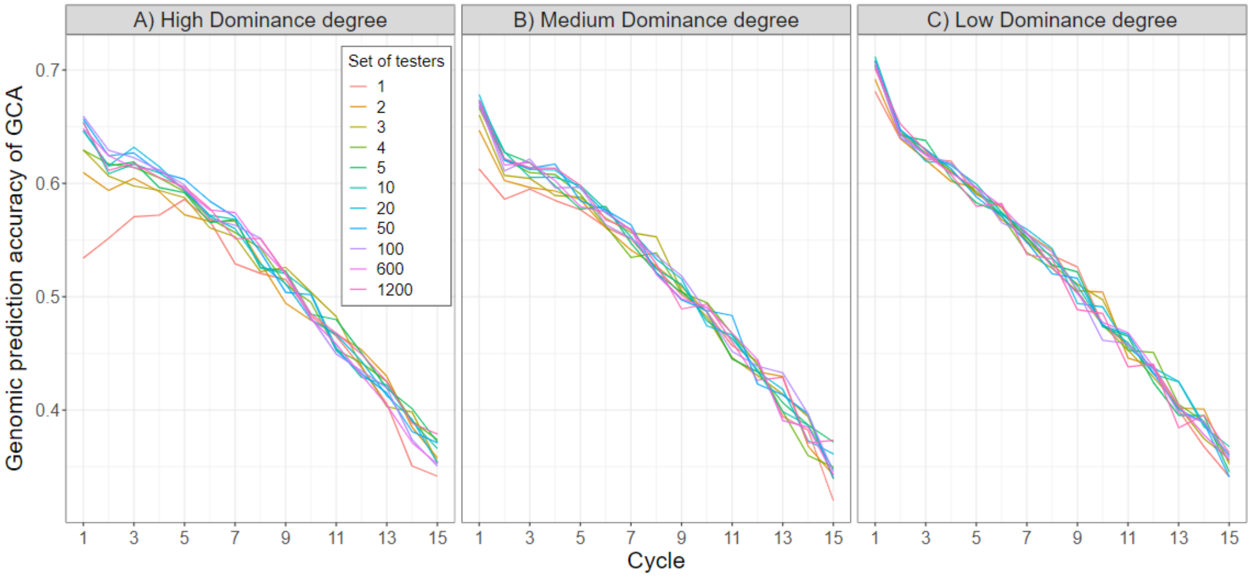


**Fig. S12** Genomic prediction accuracy of GCA by using sparse testcross designs with sets of 2, 3, 4, 5, 10, 20, 50, 100, 600 and 1200 testers, compared to a conventional testcross design with a single-tester, across 15 cycles of selection, averaged across the female and male heterotic pools and for A) high, B) medium and C) low dominance degrees. Results are averaged across all 100 simulation replications
